# High-throughput aminoacyl-tRNA synthetase engineering for genetic code expansion in yeast

**DOI:** 10.1101/2021.07.13.452272

**Authors:** Jessica T. Stieglitz, James A. Van Deventer

## Abstract

Protein expression with genetically encoded noncanonical amino acids (ncAAs) benefits a broad range of applications, from the discovery of biological therapeutics to fundamental biological studies. A major factor limiting the use of ncAAs is the lack of orthogonal translation systems (OTSs) that support efficient genetic code expansion at repurposed stop codons. Aminoacyl-tRNA synthetases (aaRSs) have been extensively evolved in *E. coli* but are not always orthogonal in eukaryotes. In this work, we use a yeast display-based ncAA incorporation reporter platform with fluorescence-activated cell sorting to screen libraries of aaRSs in high throughput for 1) incorporation of ncAAs not previously encoded in yeast; 2) improvement of the performance of an existing aaRS; 3) highly selective OTSs capable of discriminating between closely related ncAA analogs; and 4) OTSs exhibiting enhanced polyspecificity to support translation with structurally diverse sets of ncAAs. The number of previously undiscovered aaRS variants we report in this work more than doubles the total number of translationally active aaRSs available for genetic code manipulation in yeast. The success of myriad screening strategies has important implications related to the fundamental properties and evolvability of aaRSs. Furthermore, access to OTSs with diverse activities and specific or polyspecific properties is invaluable for a range of applications within chemical biology, synthetic biology, and protein engineering.

**Synopsis:** A range of flow cytometry-based screens yielded diverse translational machinery for genetic code expansion in yeast, facilitating access to new chemistries and tunable specificity profiles.

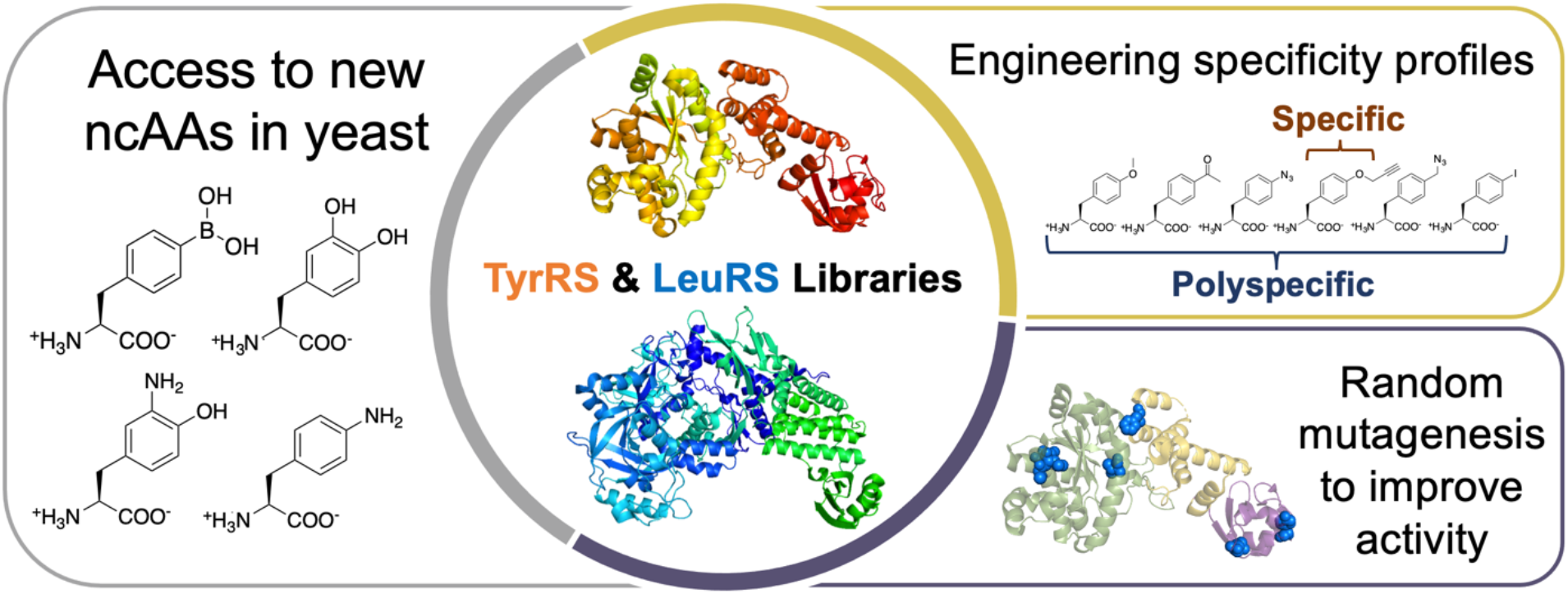

## Introduction

Diversification of protein function with genetically encoded noncanonical amino acids supports a growing number of applications in chemical and synthetic biology (ncAAs; also called unnatural amino acids, nonnatural amino acids, or nonstandard amino acids). Examples include preparation of proteins with precisely defined post-translational modifications,^*1, 2*^ dissection of protein-protein interactions,^*3, 4*^ discovery of leads for biological therapeutics,^*5, 6*^ and numerous other applications.^*7–10*^ Genetic code expansion depends on the availability of orthogonal translation systems (OTSs), comprised of aminoacyl-tRNA synthetases (aaRSs) that facilitate charging of ncAAs onto tRNAs without interfering with the host organism’s native translation machinery.^*11*^ The ncAA-charged tRNA is then translated at the ribosome in response to a stop codon or other codon that does not conventionally encode amino acids.

AaRSs have evolved to maintain the fidelity of the genetic code—that is, to precisely join canonical amino acids (cAAs) with their cognate tRNAs while discriminating against other substrates.^*12*^ Modifying the activities of aaRSs imported from other organisms to enable charging of tRNAs (usually suppressor tRNAs such as tRNA_CUA_) with ncAAs facilitates the generation of OTSs.^*11, 13*^ These orthogonal aaRS/tRNA pairs have been extensively engineered to improve the efficiency of ncAA incorporation in proteins in both prokaryotic and eukaryotic hosts, with the vast majority of efforts taking place in *Escherichia coli.^14–17^* Alteration of aaRS activities is commonly achieved using diversification of residues in the aminoacylation active site and a series of positive and negative selections to identify candidates that support incorporation of the target ncAA while discriminating against cAAs. While these approaches are powerful, the most direct and tailorable methods for the discovery and evolution of aaRSs with desirable aminoacylation activities remain unclear. For example, the extent to which aaRSs can be engineered to exhibit specificity toward a single ncAA or to enhance amino acid polyspecificity to support efficient translation with several ncAAs is not known. More broadly, there remain significant questions regarding which selection or screening strategies are most effective for discovering aaRSs that support genetic code expansion and the extent to which these strategies can be applied to genetic code expansion in organisms other than *E. coli*.

Previous work has shown that selection strategies facilitate the discovery of aaRSs capable of supporting protein translation with ncAAs. Some methods utilize life-or-death assays.^*8, 18–20*^ In yeast, a single selection strategy has been used to identify aaRS variants with improved activity toward ncAAs: *Saccharomyces cerevisiae* strain MaV203 is transformed with a plasmid encoding a GAL4 transcriptional activator gene containing two UAG codons. Suppression events drive the expression of three reporter genes (*HIS3, URA3*, and *lacZ*) that can be used to identify aaRS variants with desired readthrough properties.^*19, 20*^ This selection strategy is straightforward to implement in the laboratory and has yielded a number of aaRSs exhibiting altered amino acid substrate preferences. However, these selections tend to lead to only moderately active variants that have unknown and uncontrolled specificity profiles.^*22*^ In *E. coli*, careful adaptations of selection strategies using bioinformatics approaches^*23*^ and counterselections^*24*^ have addressed some limitations, but these approaches remain in their infancies.

Several alternative approaches to identifying aaRSs with engineered amino acid preferences have proven to be advantageous over selection-based methodologies. Again in *E. coli*, phage-assisted continuous evolution^*25*^ and several flow cytometry-based strategies have demonstrated the promise of screening approaches in discovery and engineering of aaRSs. Early efforts by Tirrell and coworkers demonstrated that aaRS activity and discrimination against cAAs can be evaluated using reporters based on GFP or on *E. coli* surface display and bioorthogonal chemistry for residue-specific^*26–29*^ ncAA incorporation. Interesting advances enhancing sitespecific ncAA incorporation in *E. coli* include high-throughput fluorescence-activated cell sorting (FACS) screens to identify highly polyspecific aaRSs^*33*^ and the use of multiplex automated genome engineering to evolve a genomically integrated aaRS for highly efficient incorporation of a target ncAA and exclusion of 237 non-target ncAAs.^*34*^ Collectively, these and other advances in screening over the past two decades have resulted in a number of aaRSs suitable for use with genetic code manipulation in *E. coli*. As a handful of OTSs in *E. coli* also exhibit orthogonality in mammalian cells, including OTSs based on pyrrolysyl-tRNA synthetases,^*35*^ some of these engineering efforts have helped diversify the genetic code expansion machinery available in higher order eukaryotes.

In contrast to *E. coli*, efforts to engineer OTSs in *S. cerevisiae* to date have used only a narrow range of approaches, despite this yeast’s critical role as both a model biological organism and as a key host in many biotechnology applications. *S. cerevisiae* is an ideal organism for evolving *E. coli* aaRSs, as OTSs based on these aaRS/tRNA pairs are naturally orthogonal and do not require the genomic modifications needed to engineer these OTSs in *E. coli* hosts.^*36*^ AaRSs evolved in yeast may also be transferrable to other eukaryotes, albeit with potential changes in activity.^*37*^ To the best of our knowledge, all yeast-based campaigns to identify aaRSs for ncAA incorporation have utilized the same set of positive and negative selections. Characterizations of selected aaRSs with advanced reporter systems indicate that the performance of aaRSs available in yeast is highly variable, and the available OTSs cover only a limited range of chemical functionalities.^*31, 38*^ Expanding the toolkit for genetic code manipulation in yeast has the potential to lead to fundamental insights into eukaryotic biology and new biotechnologies for engineering proteins and cells. Moreover, engineering OTSs in yeast may lead to a broader range of OTSs for use in other organisms that cannot currently be accessed using engineering in *E. coli*.

In this work, we established a flow cytometry-based, high-throughput screening platform for discovering aaRSs with a wide range of properties in yeast. We found that our platform supports several positive and negative screening strategies that facilitate identification of aaRSs exhibiting diverse properties, including 1) aaRSs capable of incorporating ncAAs new to yeast; 2) improved activity of an existing aaRS using random mutagenesis; 3) selective aaRSs that can discriminate between similar ncAA analogs; and 4) aaRSs that can encode structurally diverse sets of ncAAs (i.e., polyspecific aaRSs). We isolated aaRS mutants capable of supporting protein translation with numerous ncAAs, including some not previously reported to be genetically encoded in yeast, such as 3,4-dihydroxy-L-phenylalanine and 4-borono-L-phenylalanine. Screens with decreasing concentrations of ncAA in the induction media revealed variants capable of supporting moderately efficient protein translation at ncAA concentrations an order of magnitude lower than typically used in the genetic code expansion field. The expansive set of variants we isolated highlights the range of aaRS substrate preferences that can be engineered, including the discovery of highly specific EcTyrRS variants that discriminate between closely related aromatic ncAAs. On the other hand, screening for aaRSs with enhanced polyspecificity led to the isolation of many variants with broadened substrate preferences. The high plasticity of the EcLeuRS active site was especially noteworthy, with some individual variants supporting efficient translation with both aromatic and aliphatic ncAAs.

Overall, our screening platform supports the identification of aaRSs exhibiting a broad scope of properties for genetic code manipulation in yeast. When combined with the powerful protein engineering, genetic, and genomic resources available in yeast, this expanded toolkit is expected to lead to new opportunities in chemical biology, synthetic biology, protein engineering, and many other related disciplines.

## Results & Discussion

### *E. coli* aaRS saturation mutagenesis library design and construction

In order to engineer aminoacyl-tRNA synthetases (aaRSs) capable of charging diverse ncAAs to tRNA_CUA_, we constructed saturation mutagenesis libraries of the *E. coli* tyrosyl-tRNA synthetase (EcTyrRS) and *E. coli* leucyl-tRNA synthetase (EcLeuRS) (Fig. 1a–c). Previous work has shown that EcTyrRS and EcLeuRS are orthogonal to the native translation machinery present in *S. cerevisiae* and can be used to encode ncAAs in response to the amber stop codon.^*14*^ Using a combination of literature precedents and the crystal structures of the aaRSs, we identified several residues in the substrate binding pocket of each aaRS for mutagenesis.^*14*^ For the EcTyrRS library, degenerate codons at positions Y37, L71, Q179, D182, F183, L186, and Q195 were designed to reduce the size of or otherwise mutate residues in the active site that were expected to interact with the ncAA to facilitate aminoacylation of the cognate tRNA_CUA_^Tyr^ (Fig. 1, SI Tables 1 and 2). For the EcLeuRS library, positions M40, L41, S496, Y499, Y527, and H537 were randomized in addition to a T252A mutation in the editing domain known to reduce instances of tRNA_CUA_^Leu^ being charged with leucine (Fig. 1, SI Tables 3 and 4).^*39, 40*^ The permitted residues in each active site were chosen with the intention of allowing ncAAs with bulkier or longer side chains to better access aaRS substrate binding pockets to be charged to suppressor tRNAs.

**Figure 1.**
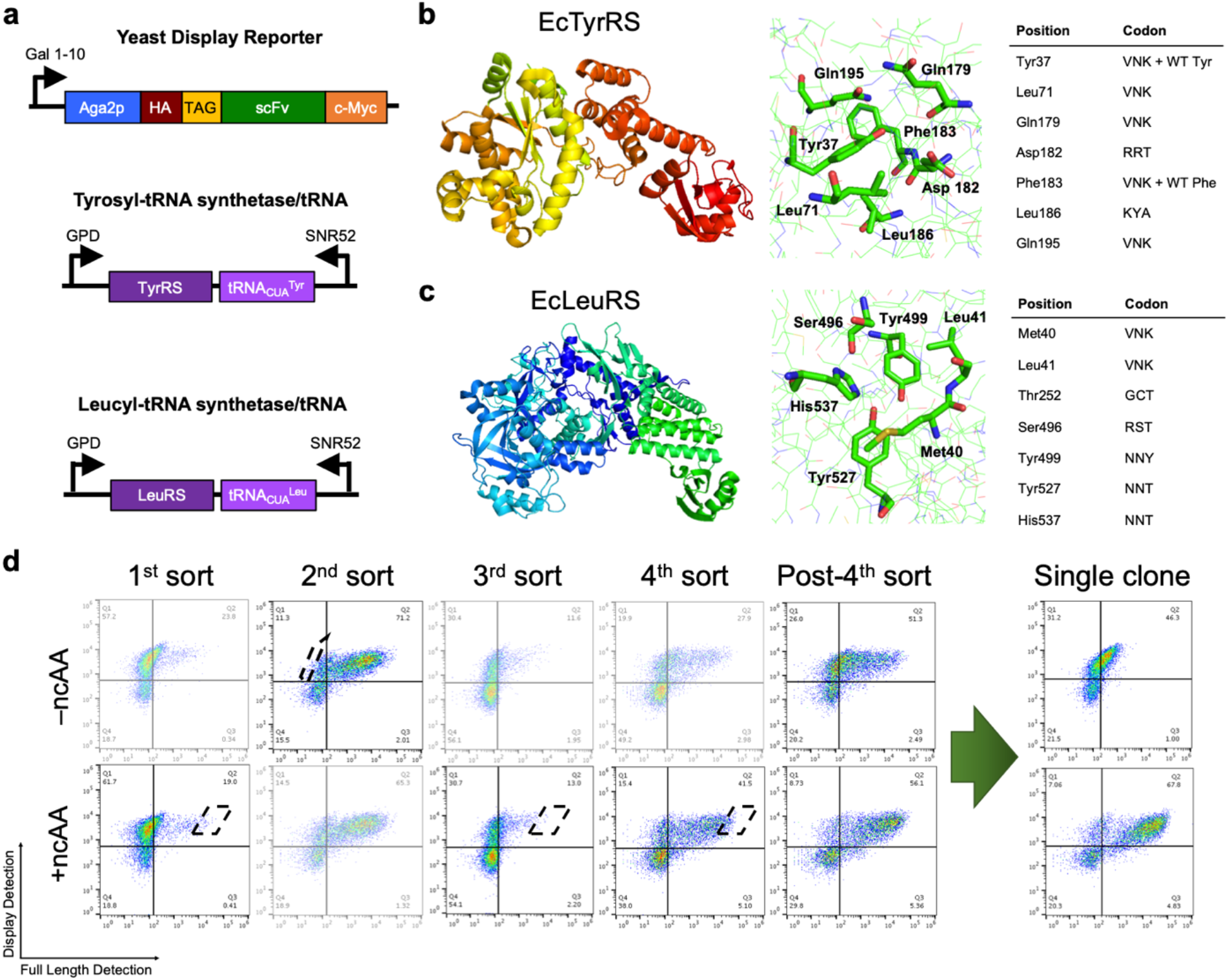
EcTyrRS and EcLeuRS library construction and FACS. **a**, Schematic representations of the genes encoding the ncAA incorporation reporter^*31*^ and the EcTyrRS and EcLeuRS with their respective tRNAs. **b**, EcTyrRS crystal structure (PDB 6HB5) with highlighted active site residues chosen for mutation. Residues and degenerate codons with library mutations are listed in the table on the right. Wildtype (WT) EcTyrRS residues were also included at positions 37 and 183. **c**, EcLeuRS crystal structure (PDB 4CQN) with highlighted active site residues chosen for mutation. Residues and degenerate codons with library mutations are listed in the table on the right. **d**, Example FACS rounds of the pooled EcTyrRS and EcLeuRS libraries for ncAA LysN3. Screens with flow cytometry-based analysis between rounds allowed for progression of screening with either a positive or negative round as needed until a population with low cAA and high ncAA incorporation was attained.

Libraries based on EcTyrRS and EcLeuRS were constructed in *S. cerevisiae* RJY100^*41*^ and evaluated for sequence diversity (SI Tables 5 and 6). Each library was co-transformed with a yeast display-based ncAA incorporation reporter that enables simultaneous detection of epitope tags located at the N- and C-termini of the reporter protein.^*31*^ The theoretical diversity of the EcTyrRS library was 1.3 × 10^8^, and the actual number of transformants was calculated to be 1 × 10^7^. Libraries containing fewer transformants than the theoretical value were considered acceptable. Although such libraries lack full mutant coverage, screening via fluorescence-activated cell sorting (FACS), with speeds generally limited to 1 × 10^8^ cells per hour, would be impractical with larger libraries in any case. All 10 randomly sequenced clones were unique. Sanger sequencing was used to evaluate all naïve and sorted aaRSs; establishing a deep sequencing platform capable of covering the full sequences of genes encoding aaRSs was beyond the scope of this work. The theoretical diversity of the EcLeuRS library was 1.9 × 10^7^, and the calculated number of transformants was 3.0 × 10^6^. Sequence characterization of the initial preparation of the EcLeuRS library revealed that out of 10 clones, the last two positions chosen for mutation (Y527 and H537) were disproportionately WT residues (SI Table 6). To correct the lack of mutations in those active site residues, a second library was constructed with slightly modified primers and was determined to contain 1.0 × 10^7^ transformants (SI Table 7). Sequence characterization of nine clones from the reconstructed EcLeuRS library revealed eight unique clones with expected mutations at all positions in the active site (SI Table 8). Since the primary goal of this work was to discover aaRSs with a range of properties, we pooled libraries encoding both EcTyrRS and EcLeuRS variants prior to sorting (SI Fig. 1). To distinguish between screens conducted with the first EcLeuRS library and the reconstructed EcLeuRS library, the original library with predominantly WT residues at positions Y527 and H537 was named Library A, and the reconstructed LeuRS library was named Library B.

### Screening aaRS libraries

Combined libraries were screened via FACS against several aromatic and aliphatic ncAA targets in search of aaRSs capable of charging these ncAAs to suppressor tRNAs (Fig. 2, SI Fig. 1). NcAA candidates exhibiting a diverse range of structures and functions were chosen as targets. This includes amino acids that have side chains that support copper-catalyzed azide-alkyne cycloaddition (CuAAC), side chains identical to naturally occurring posttranslational modifications, and side chains with potential uses in therapeutic discovery or biomaterials development. Each ncAA track was subjected to a combination of positive and negative screens. Positive sorts were conducted following induction in the presence of 1 mM ncAA and cells exhibiting high levels of full-length protein display were recovered (Fig. 1d). Negative sorts were conducted following induction in the absence of ncAAs and cells exhibiting no full-length protein display were recovered (Fig. 1d). Following several rounds of positive and negative screens, we identified enriched populations exhibiting moderate to high levels of full-length protein display following induction in the presence of ncAA and little to no full-length protein display in the absence of ncAAs (Fig. 3, SI Tables 9 and 10). Unique clones from these populations were individually evaluated for relative readthrough efficiency (RRE) and maximum misincorporation frequency (MMF)—quantitative metrics that describe the efficiency and fidelity of the aaRSs, respectively (see Supporting Information for details on RRE and MMF calculations).^*30, 31, 38*^ In general, a high RRE value corresponds to more efficient ncAA incorporation with 0 corresponding to truncation of all reporter proteins at the UAG codon and 1 corresponding to WT protein translation efficiency. MMF is a measure of the cAA misincorporation of the aaRS at the UAG codon but does not directly indicate a percentage of reporter proteins that contain a cAA at the UAG codon. Rather, MMF provides a determination of the highest possible cAA misincorporation in the worst-case scenario when there is no ncAA present.

**Figure 2.**
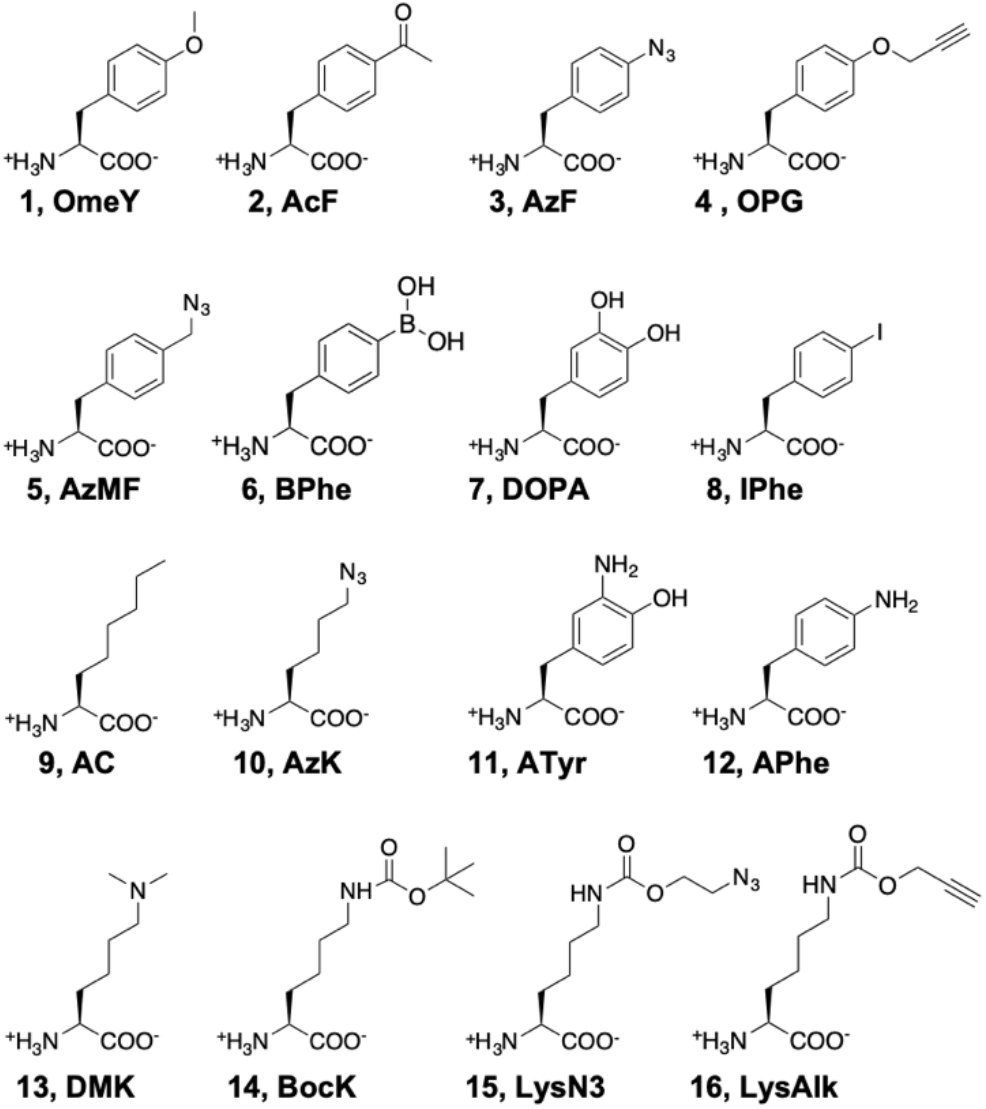
Noncanonical amino acids used in this study. **1**: *O*-methyl-L-tyrosine (OmeY); **2**: *p*-acetyl-L-phenylalanine (AcF); **3**: *p*-azido-L-phenylalanine (AzF); **4**: *p*-propargyloxy-L-phenylalanine (OPG); **5**: 4-azidomethyl-L-phenylalanine (AzMF); **6**: 4-borono-L-phenylalanine (BPhe); **7**: 3,4-dihydroxy-L-phenylalanine (DOPA); **8**: 4-iodo-L-phenylalanine (IPhe); **9**: L-α-aminocaprylic acid (AC);**10**: *N^ε^*-azido-L-lysine (AzK); **11**: 3-amino-L-tyrosine (ATyr); **12**: 4-amino-L-phenylalanine (APhe); **13**: *N^ε^, N^ε^*-dimethyl-L-lysine (DMK); **14**: Boc-L-lysine (BocK); **15**: (*S*)-2-amino-6-((2-azidoethoxy)carbonylamino)hexanoic acid (LysN3); **16**: 2-amino-6-(prop-2-ynoxycarbonylamino)hexanoic acid (LysAlk).

**Figure 3.**
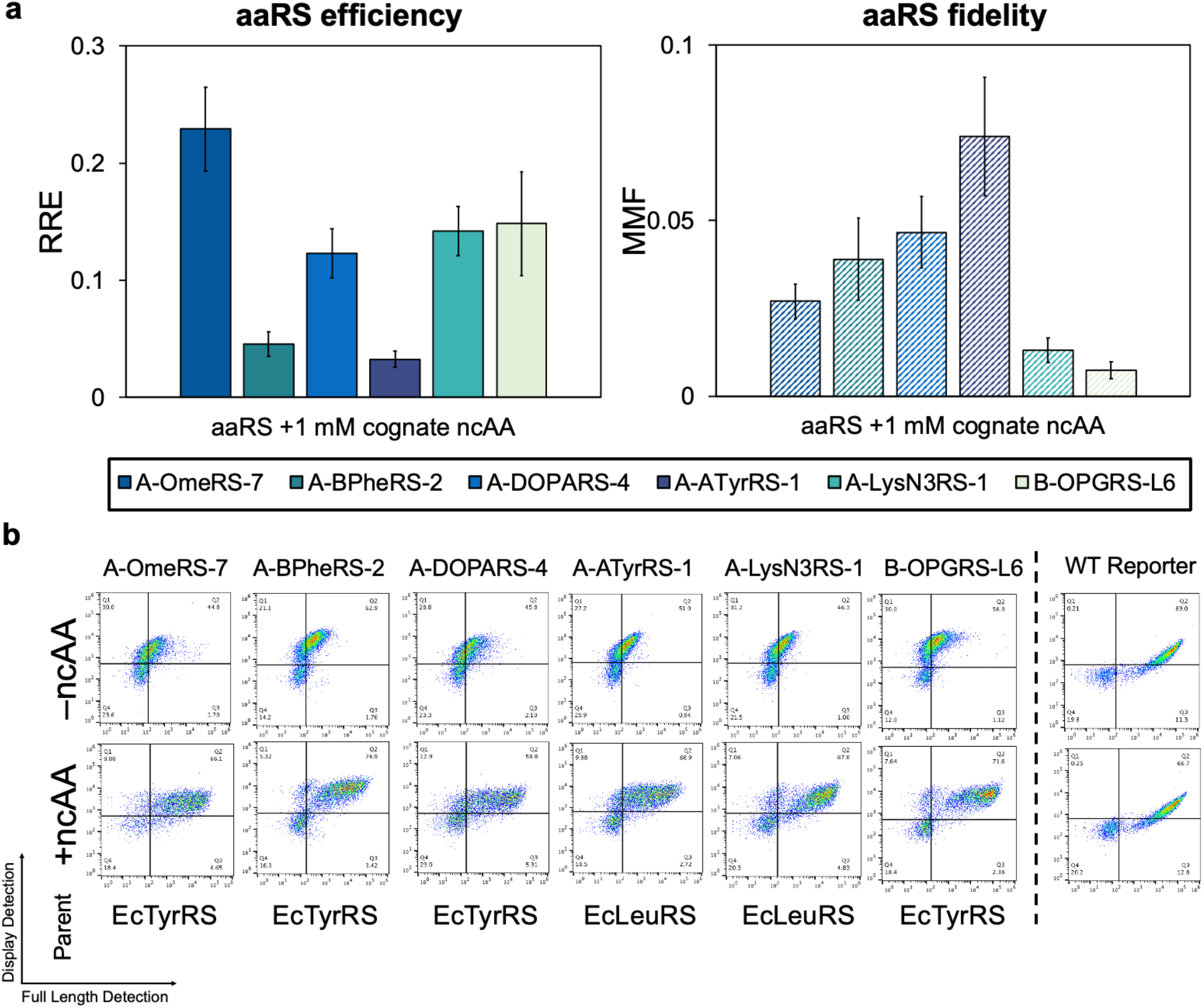
Isolated aaRSs that support translation with unique ncAAs. **a**, RRE and MMF for aaRSs that exhibit activity with target ncAAs OmeY, BPhe, DOPA, ATyr, LysN3, and OPG. **b**, Flow cytometry dot plots for each of the aaRSs in panel **a**. Only one of three biological replicates is shown for each condition (–ncAA and +ncAA). An example of WT display (no UAG codon in reporter sequence) is shown for comparison. The *E. coli* parent enzyme (EcTyrRS or EcLeuRS) is also indicated. All RRE and MMF values were derived from samples evaluated in biological triplicate. Error bars represent the standard deviation of the samples that was propagated during RRE and MMF calculations.

We isolated and characterized aaRS clones exhibiting varied translational activities with several ncAAs (Figure 3). For screens with *O*-methyl-L-tyrosine (OmeY, **1**) conducted with pooled Library A, two positive sorts and one negative sort yielded several individual clones capable of charging OmeY while discriminating against cAAs. The best clone, A-OmeRS-7, exhibited an RRE value of 0.26 ± 0.04 with an MMF value of 0.025 ± 0.004. An ncAA for which no previous studies have demonstrated translation in yeast (to the best of our knowledge) is 3,4-dihydroxy-L-phenylalanine (DOPA, **7**). We isolated five unique clones from pooled Library A that supported DOPA incorporation after four positive and one negative screens, including one (A-DOPARS-4) with an RRE value of 0.14 ± 0.025 and MMF of 0.045 ± 0.01. Screening proceeded similarly for 3-amino-L-tyrosine (ATyr, **11**), (*S*)-2-amino-6-((2-azidoethoxy)carbonylamino)hexanoic acid (LysN3, **15**), and *p*-propargyloxy-L-phenylalanine (OPG, **4**). Clones A-ATyrRS-1, A-LysN3RS-1, and B-OPGRS-L6 were isolated after five positive and two negative screens (pooled Library A), three positive and one negative screens (pooled Library A), and two positive and one negative screens (pooled Library B), respectively. One out of two unique ATyrRS clones, two out of five unique LysN3RS, and three out of four unique OPGRSs successfully charged the ncAA of interest while excluding cAAs (data not shown for all clones).

For 4-borono-L-phenylalanine (BPhe, **6**), three positive and two negative sorts of pooled Library A led to isolation of A-BPheRS-2 that was able to charge BPhe at detectable RRE levels. To our knowledge, no prior reports have demonstrated the incorporation of BPhe in proteins in yeast. Using a selection scheme in genomically engineered *E. coli*, Chatterjee and coworkers isolated an EcTyrRS variant that supports translation with BPhe in both *E. coli* and in HEK293T cells.^*36*^ While this variant exhibited some translation activity when cloned into the yeast expression system used in this work, it appears to be less efficient than A-BPheRS-2 (SI Fig. 2). Interestingly, three out of five unique APheRS clones from pooled library A sorts showed some 4-amino-L-phenylalanine (APhe, **12**) incorporation and no cAA misincorporation in the qualitative flow cytometry dot plots, but corresponding RRE values were calculated to be zero (SI Fig. 3). This confirms that our screening approach supports enrichment of clones exhibiting weak translational activity, but also suggests that quantitative assessments of ncAA incorporation with the stringent RRE metric may not fully capture aaRS activity levels in cases where partial readthrough is evident.

To further characterize the ncAA-containing proteins, several promising aaRS mutants were co-transformed with a secreted reporter protein, purified using Protein A column chromatography, and evaluated via MALDI mass spectrometry analysis. Masses for each ncAA-containing peptide were consistent with the expected masses (SI Fig. 4, SI Table 11). Taken as a whole, flow cytometry analysis and mass spectrometry characterizations confirm that the aaRS libraries screened here contain clones that support the incorporation of a variety of ncAAs into proteins in yeast. This validates our FACS-based discovery platform in yeast and provides opportunities to more systematically determine the range of chemically diverse ncAAs that EcTyrRS and EcLeuRS are capable of charging to suppressor tRNAs to support protein translation.

### EpPCR library construction and screening

While we successfully isolated aaRSs capable of charging new ncAAs from the EcTyrRS and EcLeuRS libraries during initial sorts, some clones exhibited only modest stop codon readthrough. We sought to further improve the activity of one of these aaRSs with the use of random mutagenesis mediated by error-prone PCR (epPCR). Some prior work has demonstrated the utility of epPCR in enhancing aaRS activity in *E. coli.^43–48^* However, this approach has never been explored in yeast, and, more broadly, this approach remains underexplored in aaRS engineering. In this case, an EcTyrRS mutant that charges DOPA, A-DOPARS-8, was PCR-amplified with two concentrations of mutagenic dNTPs to introduce random point mutations. A DOPARS parent was chosen over other potential candidates, such as an ATyrRS or BPheRS, due to the potential utility of DOPA in biomaterials and bioadhesive applications, among other uses. Two epPCR libraries, DOPARS-1X and DOPARS-5X, were constructed in *S. cerevisiae* RJY100 with 1.6 × 10^7^ transformants for DOPARS-1X and 2.0 × 10^7^ transformants for DOPARS-5X. Based on sequence characterization of 12 clones per library, the average number of point mutations per gene in DOPARS-1X was determined to be 4, and in DOPARS-5X the average number of mutations per gene was determined to be 17; additionally, all clones sequenced from the naïve libraries were unique.

Both naïve libraries were pooled and screened via FACS following reporter induction in the presence of 1 mM DOPA. Inductions in subsequent positive screening rounds utilized progressively lower concentrations of DOPA (0.1 and 0.05 mM). After four positive screens and one negative screen, characterization of individual clones revealed several variants with improved ability to support protein display following induction in the presence of 0.1 mM or 1 mM DOPA in comparison to the parent clone (Fig. 4, SI Table 12, SI Fig. 5). It is worth noting that based on the average number of mutations in the clones that were enriched via FACS, these aaRSs were likely isolated from the DOPARS-1X library; the significantly higher numbers of mutations present in clones from the DOPARS-5X library were not beneficial in this case. Regardless, given that most studies with aaRSs utilize ncAA concentrations of 1 mM or higher, the substantial improvement of activity at 0.1 mM is noteworthy.

**Figure 4.**
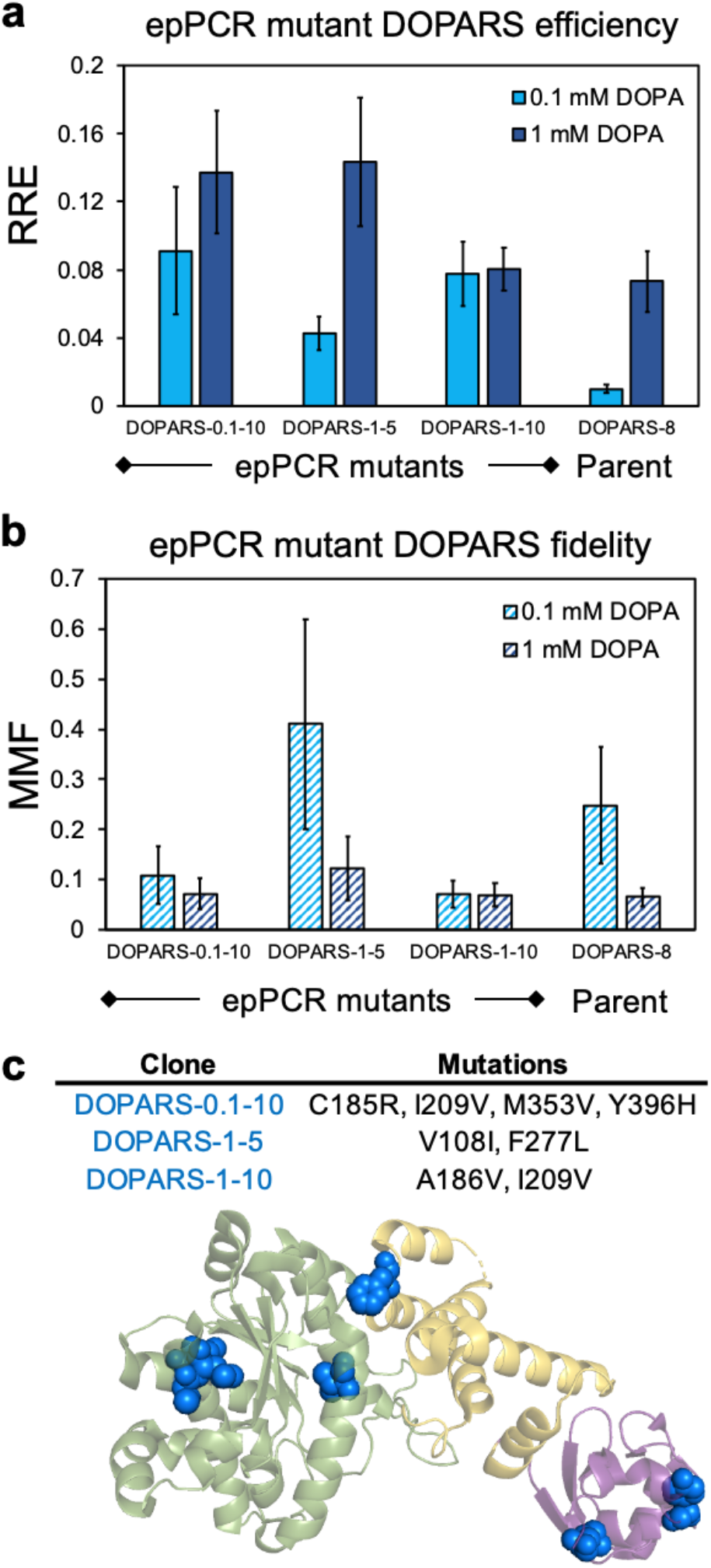
Performance of aaRS mutants isolated from random mutagenesis libraries. **a**, RRE of three epPCR mutants that outperform the parent DOPARS at 0.1 mM, 1 mM, or both concentrations of DOPA. **b**, MMF of three epPCR mutants exhibiting comparable fidelities to that of the parent DOPARS at both 0.1 and 1 mM DOPA. **c**, Crystal structure of EcTyrRS with mutated residues identified following random mutagenesis and screening highlighted in blue. The N-terminal catalytic domain is shown in green, the anticodon recognition domain is shown in yellow, and the C-terminal domain is shown in purple. The crystal structure in this figure was derived from PDB 6HB5. All RRE and MMF values were derived from samples in biological triplicate and error bars represent the standard deviation of the samples that was propagated during RRE and MMF calculations.

Many unique epPCR DOPARSs were identified from sequence characterization, with no clear enrichment for one particular sequence. However, several trends emerged regarding the positions of mutations. Most commonly, clones contained mutations directly adjacent to the active site or in the tRNA binding domain (Fig. 4c, SI Table 12). Clones DOPARS-0.1-2, −5, −9, −10 and DOPARS-1-7, −8, −9, −10, −11, and −12 all contained one or more of the following mutations in the active site: Q18R, T37G, M179N, C185Y/R, A186T/V, or I209V (based on the sequence of A-DOPARS-8). In total, half of the epPCR-derived mutants sequenced had one or more mutations in the active site. Additionally, many variants contained mutations in the tRNA^Tyr^ recognition domain. Key residues in the EcTyrRS tRNA binding domain have been previously reported elsewhere,^*49*^ and recently an improved DOPARS variant of the *Methanocaldococcus jannaschii* TyrRS possessing mutations in the anticodon binding domain was reported by Ellington and coworkers.^*44*^ One of the epPCR clones isolated here, DOPARS-0.1-9, had a mutation at a known tRNA recognition site: K377E. Several additional clones had mutations that may be involved with tRNA recognition based on previous structural characterizations, but these mutations are not at locations where direct effects on tRNA recognition have been experimentally demonstrated.^*50, 51*^ While these mutations are not located in the active site or at known tRNA recognition sites, their occurrence in multiple unique clones suggests that they may be important for enhancing EcTyrRS activity and thus merit further study in future work. In any case, our findings here indicate that our FACS-based screen supports increasingly stringent screens following epPCR to identify variants with enhanced readthrough activity, even at reduced ncAA concentrations.

### Modified screens for isolation of aaRSs with desired specificity profiles for OPG incorporation

Having validated positive and negative screens with our FACS-based approach, we sought to investigate alternative induction conditions prior to FACS to isolate highly specific aaRSs—aaRSs that charge a single ncAA to the cognate tRNA_CUA_ and do not mischarge structurally related ncAAs. Previous selections in yeast^*19, 52–54*^ and the sorts described above did not attempt to control ncAA substrate specificity preferences other than the preference for the target; such conditions can lead to polyspecific aaRSs.^*33, 55, 56*^ For these experiments, we used a group of six structurally similar aromatic ncAAs: OmeY, *p*-acetyl-L-phenylalanine (AcF, **2**), *p*-azido-L-phenylalanine (AzF, **3**), OPG, 4-azidomethyl-L-phenylalanine (AzMF, **5**), and 4-iodo-L-phenylalanine (IPhe, **8**). Based on our previous characterizations of EcTyrRS variant TyrAcFRS^*57*^ and EcLeuRS variant LeuOmeRS,^*58*^ we expected that similarities between these ncAAs would make it difficult for aaRS variants to selectively charge only a single ncAA without also charging others.^*31*^

We investigated the possibility of isolating aaRSs capable of charging OPG but not the other five ncAAs of this set of six ncAAs with our FACS-based screens. Starting from the pooled EcTyRS and EcLeuRS B libraries, we modified the negative screens and added 1 mM of all five non-target ncAAs during induction, while standard positive screens remain unchanged (1 mM OPG was added during induction for positive sorts). After two positive and three negative rounds of screening, RRE and MMF evaluation of individual clones isolated from these specificity sorts reveal several clones that exhibit substantial translational activity with OPG but nearly undetectable levels of translation with the other ncAAs in the set (Fig. 5a). Three unique clones, SpecOPGRS-3, −7, and −9, were able to charge OPG at RRE values of 0.05–0.20, with RRE <0.01 for each of OmeY, AcF, AzF, AzMF, and IPhe. In contrast, clone B-OPGRS-L6, which was isolated using conventional positive and negative (no off-target ncAAs) screens, exhibited high levels of polyspecificity, supporting protein translation with four off-target ncAAs (AcF, AzF, OmeY, and IPhe) (Fig. 6a, SI Fig. 6a). Out of the four unique OPGRS clones from conventional sorts only three were active and sequence similarity between these clones and clones isolated from polyspecificity sorts (see below) suggested that the OPGRSs would non-specifically encode ncAAs other than OPG. This was confirmed for B-OPGRS-L6 but not for the other two active, unique OPGRSs. There are marked differences between the specificity profiles of aaRSs isolated using specificity screening methods and aaRSs isolated using more standard screening approaches. Tailoring evolutionary pressures can drastically improve the substrate specificity of an aaRS isolated during screens.

**Figure 5.**
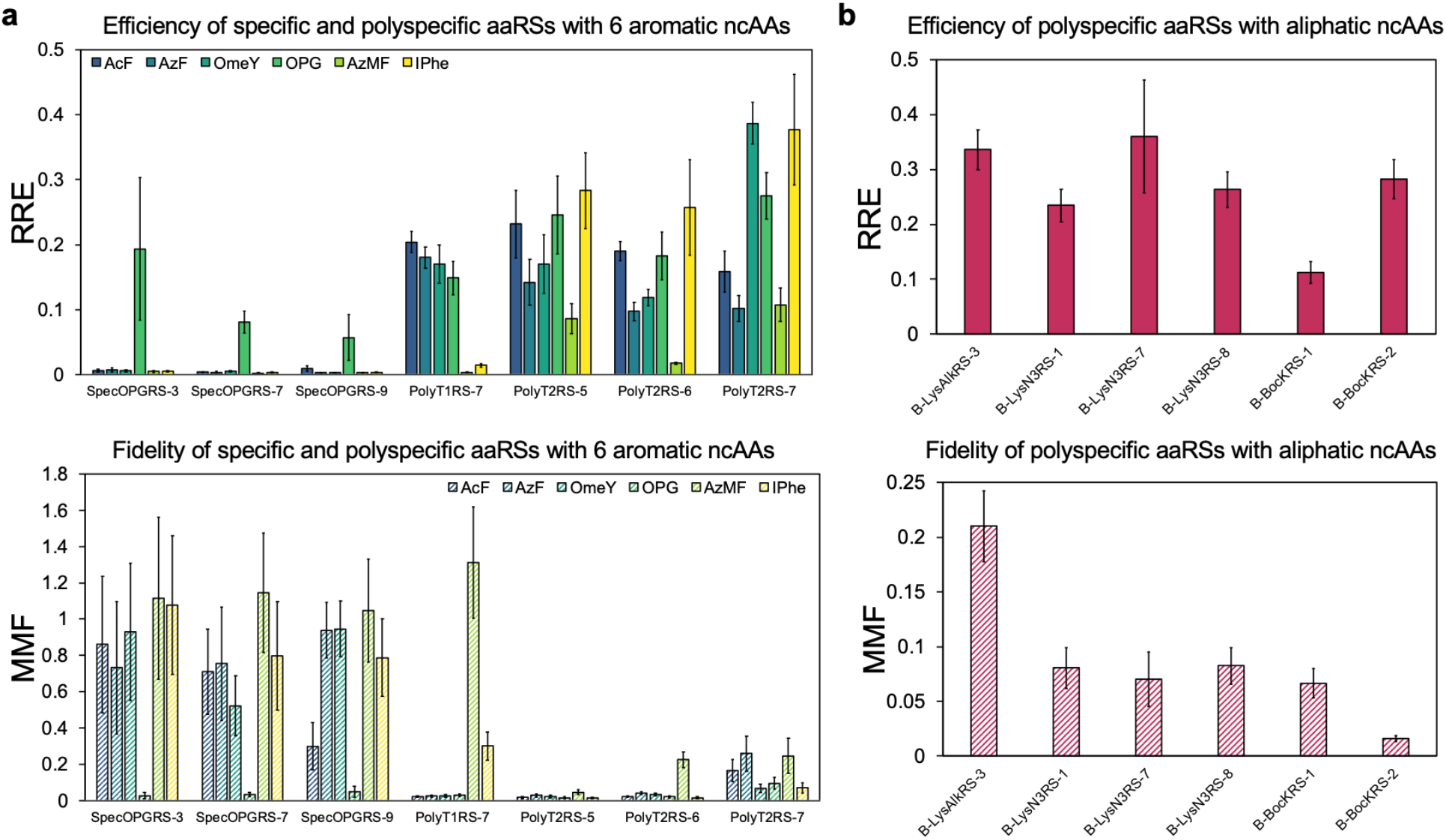
NcAA incorporation efficiency and fidelity of specific and polyspecific aaRSs isolated via FACS. **a**, Evaluation of RRE and MMF of three specific aaRSs (SpecOPGRS-3, SpecOPGRS-7, and SpecOPGRS-9) and four polyspecific aaRSs (PolyT1RS-7, PolyT2RS-5, PolyT2RS-6, and PolyT2RS-7) from Track 1 and Track 2 polyspecificity sorts with six aromatic ncAAs used during screening. For Track 1 sorts, all six ncAAs were added during the induction step prior to positive sorts. For Track 2 sorts, only one ncAA was added during induction prior to a positive sort and subsequent positive sorts were performed with a distinct ncAA from the group of six. **b**, Evaluation of RRE and MMF of six aaRSs originating from a Track 1 polyspecificity screen that encode aliphatic ncAAs LysAlk, LysN3, and BocK. AaRSs were evaluated here for translation activity with the ncAA included in the name of the clone (e.g., B-LysAlkRS-3 was evaluated for translation activity with LysAlk). The concentration of ncAA used during induction was 1 mM for all samples in this figure. All RRE and MMF values were derived from samples evaluated in biological triplicate with error bars representing the standard deviation of the samples that was propagated during RRE and MMF calculations.

**Figure 6.**
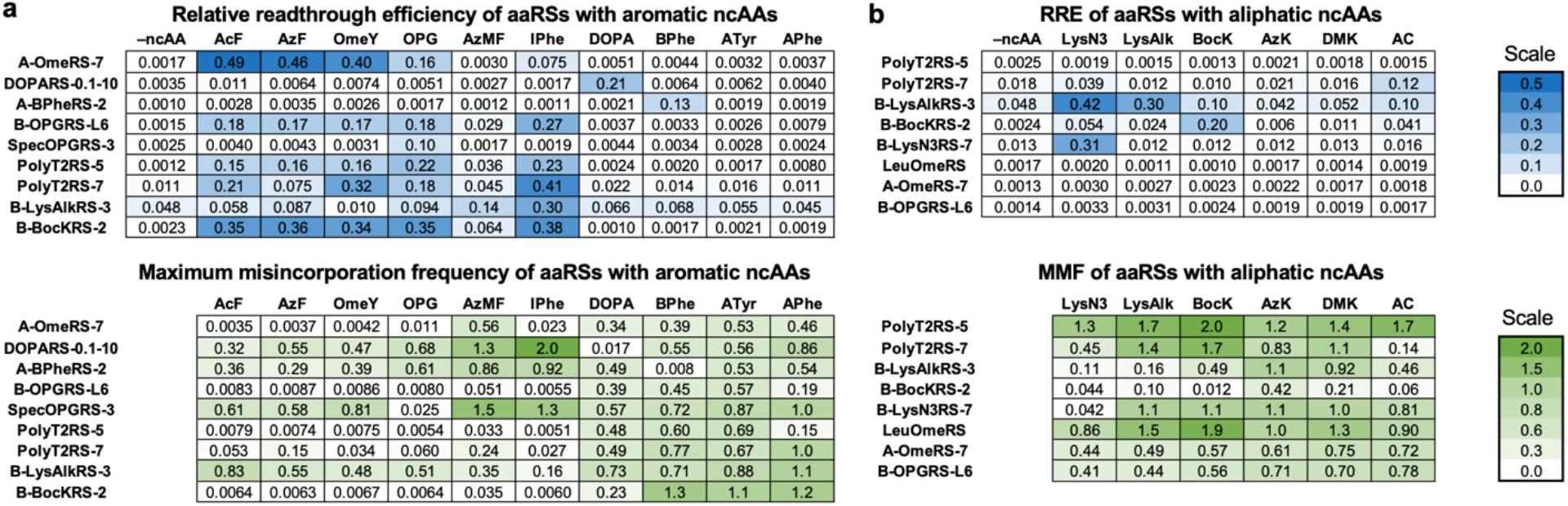
Comprehensive evaluation of aaRS activity with aromatic and aliphatic ncAAs. **a**, RRE and MMF for nine aaRSs with 10 aromatic ncAAs. **b**, RRE and MMF for eight aaRSs with six aliphatic ncAAs. The concentration of ncAA used during induction was 1 mM for all samples in this figure. All RRE and MMF values were derived from samples evaluated in biological triplicate and error can be found in SI Figure 6.

### Modified screens for isolation of aaRSs with desired polyspecificity profiles

We also sought to investigate whether we could enhance aaRS polyspecificity using altered screening methods via two strategies: Track 1 and Track 2. For Track 1, we added all six of the same group of aromatic ncAAs (OmeY, AcF, AzF, OPG, AzMF, and IPhe) at 1 mM to the induction media for positive sorts. In a Track 1 series of screens consisting of three consecutive positive sorts followed by a single negative sort, characterization yielded four unique EcTyrRS mutants and one EcLeuRS mutant (SI Tables 9 and 10). For Track 2, only one of the six aromatic ncAAs was added during induction per positive sort round, with a different ncAA added for each subsequent positive sort. The B pooled libraries were screened first using AzF, followed by a positive screen with AcF, a negative screen, and two consecutive positive screens for AzMF due to our observation of lower incorporation of AzMF in comparison to the other five ncAAs during flow cytometry characterization of the sorted populations (SI Fig. 7). Out of 11 individual clones isolated from the Track 2 sorted library population, seven were unique: five EcTyrRS variants and two EcLeuRS variants (SI Tables 9 and 10).

Unique clones from Track 1 and 2 polyspecificity sorts were first evaluated for incorporation of each of the six aromatic ncAAs and misincorporation of cAAs. Evaluation of the efficiency (RRE) and fidelity (MMF) of the most active polyspecific aaRSs demonstrates that the aaRSs were able to encode several of the group of six ncAAs and also reveals a key difference in outcome between the two sort tracks (Fig. 5a). For Track 2 sorts, we performed two consecutive AzMF positive sorts to ultimately isolate aaRSs that encoded all six ncAAs, as flow cytometry characterizations between sorts indicated reduced AzMF incorporation compared to the other five ncAAs. In the case of Track 1, the most active clone (PolyT1RS-7, an EcLeuRS variant) encodes only four out of the six ncAAs, which was known during flow cytometry characterizations between sort rounds but was not controllable since all six ncAAs were added during all inductions. Interestingly, an identical sequence to known polyspecific enzyme TyrAcFRS^*31, 57*^ was found seven times in the Track 1 population and two times in the Track 2 population. The only positions that differed between the polyspecific aaRSs derived from EcTyrRS are at positions Y37, L71, and V72. Position V72 was not included in the original set of active site residues chosen for mutation and did not appear in the initial EcTyrRS library characterization. However, mutations at V72 appeared in several EcTyrRS clones isolated from different screens and may be attributable to a mutation introduced during PCR amplification prior to library construction.

Each polyspecificity sort strategy has its advantages and shortcomings. Track 1 is straightforward but does not provide control over which ncAA is being encoded by the aaRSs for any given sort round. Track 2 requires careful monitoring of individual ncAA incorporation by the sorted library populations after each sort but allows for more control of individual ncAA incorporation as needed. By making simple modifications to the induction step prior to screening, aaRSs with polyspecific characteristics can be isolated for use in applications where a single aaRS that can encode multiple ncAAs is desirable. Furthermore, both strategies could be readily applied to other types of screens for isolation of aaRSs with tailored specificity profiles in *E. coli* or other organisms, and with screens that use methods other than FACS.

### Characterizing polyspecific aaRSs for activity with aliphatic ncAAs

To further evaluate the extent of polyspecificity of the final enriched Track 1 library population, we tested the population for incorporation of 21 ncAAs (SI Fig. 8). Interestingly, while the Track 1 aaRSs were only screened against a group of aromatic ncAAs, they also appeared to support translation with several aliphatic ncAAs at low levels. To identify aaRSs from the Track 1 library population exhibiting polyspecificity for ncAAs beyond the original six aromatic ncAAs, we further screened the library against three aliphatic ncAAs: Boc-L-lysine (BocK, **14**), LysN3, and 2-amino-6-(prop-2-ynoxycarbonylamino)hexanoic acid (LysAlk, **16**). A single positive screen led to the isolation of a number of EcLeuRS variants capable of supporting efficient incorporation of one or more of these ncAAs into proteins (SI Tables 9 and 10). Some major trends included the isolation of one particularly active, polyspecific clone 11 times out of 32 colonies sequenced across the three sorts, and frequent amino acid mutations to glycine at both positions S496 and H537. Convergence to H537G has been observed in previous selections with EcLeuRS.^*59*^ As S496 has not, to our knowledge, been included in EcLeuRS saturation mutagenesis library design, convergence to S496G is particularly intriguing.

We evaluated the efficiency and fidelity of ncAA incorporation for several clones from each population and discovered that some were able to encode the aliphatic ncAA target at levels comparable to the Track 1 and Track 2 RRE values for the original six aromatic ncAAs with similarly low misincorporation of cAAs (Fig. 5b). It is noteworthy that while very few EcLeuRS variants were identified during characterization of the original Track 1 and 2 screens, all the aaRSs isolated from the aliphatic ncAA sorts were derived from EcLeuRS. Overall, these results demonstrate that polyspecificity screens can lead to the identification of aaRSs that support translational activity with structurally diverse ncAAs spanning aromatic and aliphatic side chains.

### Comparison between aaRSs sorted using conventional versus modified methods

To directly compare the performance of aaRSs isolated using distinct screening methods, we determined the efficiency and fidelity of nine aaRSs with 10 aromatic ncAAs and eight aaRSs with six aliphatic ncAAs (Fig. 6). For aaRSs evolved using conventional positive and negative screens, clones exhibited variable levels of specificity or polyspecificity. For example, A-BPheRS-2 and DOPARS-0.1-10 both exhibited specificity towards the ncAAs BPhe and DOPA, respectively, whereas A-OmeRS-7 and B-OPGRS-L6 were able to encode five of the group of 10 aromatic ncAAs at moderate to high levels (Fig. 6a, SI Fig. 6a). For aaRSs isolated from intentionally polyspecific or specific sorts, the behavior of the aaRSs were consistent with types of screens employed. Specificity clone SpecOPGRS-3 only supports stop codon suppression with OPG and not any of the other aromatic ncAAs evaluated here. Polyspecificity clones PolyT2RS-5 (EcTyrRS variant) and PolyT2RS-7 (EcLeuRS variant) perform well with five out of the original six aromatic ncAAs they were screened for incorporation with and support translation with the sixth ncAA, AzMF, at low but detectable levels (AzMF incorporation is apparent in the flow cytometry dot plots, but here the stringency of the RRE metric resulted in the calculation of background-level values; see SI Fig. 3 for dot plots). Interestingly, these polyspecificity clones do not appear to support high levels of incorporation with any of the other four aromatic ncAAs tested here.

The two aaRSs originally identified using Track 1 polyspecific methods and then isolated after subsequent screens with aliphatic ncAAs LysAlk and BocK both exhibit highly polyspecific behavior toward the original set of six aromatic ncAAs, with notably lower AzMF incorporation. We note that PolyT2RS-5 also supports some degree of translation with APhe based on flow cytometry dot plots, although again RRE calculations resulted in values close to background levels (SI Fig. 3). We attribute the low RRE values that occur, despite the presence of detectable readthrough events in flow cytometry plots, to the following: cells that have lost the suppression machinery (due to plasmid loss) and are incapable of reading through the UAG codon are included in the determinations of N- and C-terminal display levels, which lowers the overall median fluorescence intensity (MFI). In other words, cells exhibiting zero readthrough lower the median fluorescence of populations that exhibit low levels of readthrough, obscuring the detected activity. With enough reduction, the result is near-zero RRE values that do not capture the readthrough events that are apparent in flow cytometry plots. This effect is most noticeable for aaRSs exhibiting highly variable cell-to-cell activity levels or generally low activity. Despite near-zero RRE values, corresponding MMF values were still low in comparison to aaRSs with no qualitative readthrough and zero RRE values. This raises the possibility that low MMF values may be a way to identify translational activity in cases of low or variable translational activity, but further investigations are needed to thoroughly investigate this conjecture. In addition, modifications to gating strategies or the use of mean or geometric mean fluorescence intensities in place of median fluorescence intensities for calculating RRE may lead to values that better capture low but nonzero aaRS activity levels.

We also evaluated several aaRSs against a panel of aliphatic ncAAs (Fig. 6b, SI Fig. 6b). Polyspecificity clone PolyT2RS-5 showed virtually no incorporation of any of the aliphatic ncAAs tested, but EcLeuRS variant PolyT2RS-7 incorporated L-α-aminocaprylic acid (AC, **9**) with an RRE value of 0.12 ± 0.025. LeuOmeRS, A-OmeRS-7, and B-OPGRS-L6, all of which were isolated following screens or selections for incorporation of a single ncAA, did not support stop codon readthrough with any of the aliphatic ncAAs tested. The highest performing aliphatic ncAA-encoding aaRSs (that were screened for aromatic ncAA incorporation prior to screens for incorporation of individual aliphatic ncAAs) were B-LysAlkRS-3, B-BocKRS-2, and B-LysN3RS-7. B-LysN3RS-7 encoded LysN3 efficiently with an RRE value of 0.31 ± 0.051 but did not encode any other aliphatic ncAAs tested. B-BocKRS-2 similarly encoded BocK well (RRE of 0.20 ± 0.033) but showed little to no activity for any other aliphatic ncAAs. Unlike its BocK and LysN3 counterparts, B-LysAlkRS-3 not only supported translation with LysAlk efficiently (RRE of 0.30 ± 0.024), but also was translationally active with LysN3, BocK, and AC. Notably, B-LysAlkRS-3 exhibits a slightly higher background level of cAA misincorporation, as determined by MMF measurements. The ability to take a polyspecificity screen designed to encode aromatic ncAAs and isolate aaRSs additionally capable of charging aliphatic ncAAs further demonstrates the extent of polyspecificity achievable using slightly modified induction steps prior to aaRS screens.

Another interesting trend we observed was differences in polyspecificity between EcTyrRS and EcLeuRS variants isolated from various screens. The highest performing EcTyrRS variants isolated from screens for aminoacylation of OmeY and OPG exhibited polyspecificity with aromatic ncAAs AcF, AzF, OmeY, OPG, and IPhe (as well as AzMF in the case of B-OPGRS-L6), but these variants were not able to support translation with any of the aliphatic ncAAs tested (Fig. 6). On the other hand, some EcTyrRS clones such as DOPARS-0.1-10 and A-BPheRS-2 showed excellent substrate specificity for their cognate ncAAs. While PolyT2RS-5 efficiently produced proteins containing five out of the six aromatic ncAAs it was originally screened against, and demonstrated detectable translational activity with APhe, it was not able to charge the other three aromatic or six aliphatic ncAAs evaluated. However, EcLeuRS clone PolyT2RS-7 was able to support translation with the same aromatic ncAAs as PolyT2RS-5 as well as LysN3 and AC at low but detectable levels. Similarly, EcLeuRS clones LysAlkRS-3 and BocKRS-2 both supported stop codon suppression with some aromatic ncAAs, as well as multiple aliphatic ncAAs. In particular, LysAlkRS-3 charged BocK and AC at moderate levels and LysAlk and LysN3 at high levels. These results suggest that EcLeuRS variants are capable of exhibiting broader levels of polyspecificity than EcTyrRS variants. Taken together, the EcTyrRS and EcLeuRS variants isolated in this work exhibit a range of (poly)specificity profiles, some of which were only accessible after tailoring screening conditions to maximize (poly)specificity. These findings raise interesting questions about the limits of aaRS substrate specificity and polyspecificity and point to strategies for systematically investigating such questions.

## Conclusions

In this work, we employed flow cytometry-based screens to discover aaRSs exhibiting a wide range of properties for genetic code expansion in yeast. To the best of our knowledge, this is the first report utilizing such approaches in yeast to engineer OTSs. Isolation of clones from saturation mutagenesis EcTyrRS and EcLeuRS libraries led to numerous variants supporting protein translation with a broad set of ncAAs, including DOPA, BPhe, and other ncAAs that had not previously been genetically encoded in yeast. However, we note that not all ncAAs we attempted discovery with yielded functional aaRSs, suggesting that further library design based on our findings may be necessary to expand the substrates that can be accepted by engineered EcTyRSs and EcLeuRSs. With this platform, we were also able to improve the properties of an existing aaRS via error-prone PCR. Random mutagenesis of a DOPARS variant followed by increasingly stringent screening led to identification of clones that support improved protein translation with DOPA even at reduced ncAA concentrations. The facile discovery of improved variants in a single round of mutagenesis suggests the possibility that this platform will facilitate more extensive, multi-round aaRS discovery and mutagenesis campaigns in the future—a generally underexplored route to modifying aaRS activity. Moreover, these findings highlight the strong potential to enhance OTS performance by broadly exploring aaRS diversification strategies beyond aaRS aminoacylation active sites; this observation is consistent with the findings of other recent work in this area.^*25, 43–45*^

By tailoring the conditions under which positive and negative screens for ncAA incorporation were performed, we were able to isolate aaRSs with a wide range of specificity and polyspecificity profiles. Inclusion of off-target ncAAs during negative screens enabled us to identify aaRS variants that incorporate OPG into proteins while discriminating against five closely related aromatic ncAAs. In the absence of these ncAAs, we were only able to isolate polyspecific aaRSs that support translation with OPG, in line with previous reports.^*53*^ On the other hand, when attempting to identify clones exhibiting higher levels of polyspecificity, both induction in the presence of multiple ncAAs (Track 1) and induction in the presence of different ncAAs in subsequent FACS rounds (Track 2) led to the isolation of clones exhibiting enhanced polyspecificity. While each of these approaches was successful, we note some important differences between the approaches. Track 1 is straightforward because the same set of ncAAs is utilized during induction for each positive screen, but this approach does not allow for control over which individual ncAAs are incorporated into reporter proteins during screening. Track 2 requires careful (and somewhat laborious) monitoring of individual ncAA incorporation after each sort, but these evaluations allow for the inductions prior to each sort to be precisely modified to maximize polyspecificity across a set of ncAAs. Intriguingly, by taking a population already enriched for polyspecific aromatic ncAA incorporation and screening against a set of aliphatic ncAAs, we identified aaRSs capable of supporting efficient protein translation with both aromatic and aliphatic ncAAs. Access to both highly specific aaRSs and highly polyspecific aaRSs are both desirable in different applications of genetic code manipulation.

Our powerful screening platform for engineering aaRSs in yeast generated many mutants capable of incorporating structurally diverse ncAAs into proteins. The breadth of aaRS properties accessed in this work underscores the excellent plasticity and “evolvability” of these enzymes. We attribute the functional diversity of the mutants reported here primarily to the carefully controlled screening conditions, with both flow cytometry gating strategies and well-defined induction conditions playing key roles in biasing screening outcomes. There are several ways in which the findings described here could be extended further in future work. First, detailed sequence-activity relationships for the aaRSs investigated here may be attainable, especially if deep sequencing methodologies can be applied.^*23*^ Second, understanding the relationship between the observed translation properties reported here and underlying OTS properties (e.g., kinetic constants of aaRSs, expression levels of OTS components, and expression conditions) could lead to better understanding of how best to efficiently prepare ncAA-containing proteins in high yields and purities. AaRS kinetics have the potential to provide critical insights into how best to enhance the properties of the protein translation machinery used to manipulate the genetic code. Third, the availability of highly specific OTSs has the potential to facilitate genetic code expansion to include multiple ncAAs in the same protein, even when the two ncAAs of interest are similar in structure. Finally, polyspecific aaRSs have potential utility in applications where protein medicinal chemistry to systematically explore the effects of different ncAA side chains on protein properties is desirable.^*60, 61*^ Our findings begin to explore the most efficient way to select sets of ncAAs that lead to tightly controlled aaRS activity using modified specificity profile screens. Additional work using various sets of ncAAs may lead to maximally polyspecific aaRSs that are still able to discriminate against cAAs.

Beyond changing the ncAA substrate preferences of aaRSs, our findings indicate that there remain substantial opportunities to enhance aaRS activities by exploring mutations beyond aminoacylation active sites. Here, a single round of random mutagenesis led to the discovery of aaRS variants possessing a diverse set of mutations exhibiting improved translational activity. These findings are consistent with screens performed by other groups in *E. coli*, where random mutagenesis and screening led to rapid improvement of aaRS properties by the introduction of mutations at positions outside of aminoacylation active sites. Further studies using random mutagenesis, deep mutational scanning,^*62*^ or other approaches to facilitate more comprehensive exploration of the sequence spaces surrounding known aaRSs represents a major opportunity for understanding and engineering aaRSs. In addition to advancing discovery of highly active, next-generation OTSs, such studies are expected to lead to fundamental insights into the biochemical properties of aaRSs. Overall, the availability of a high-throughput screening platform for aaRSs in yeast broadens opportunities for generating versatile aaRSs suitable for use in genetic code manipulation in yeast, mammalian cells, and other eukaryotes. Such tools are expected to facilitate dissection of basic biological and biochemical phenomena as well as myriad applications at the interface of chemical biology, synthetic biology, and protein engineering.

## Materials and Methods

All restriction enzymes used for molecular biology were from New England Biolabs (NEB). Synthetic oligonucleotides for cloning and sequencing were purchased from Eurofins Genomics or GENEWIZ. All sequencing in this work was performed by Eurofins Genomics (Louisville, KY) or Quintara Biosciences (Cambridge, MA). Epoch Life Science GenCatch™ Plasmid DNA Mini-Prep Kits were used for plasmid DNA purification from *E. coli*. Yeast chemically competent cells and subsequent transformations were prepared using Zymo Research Frozen-EZ Yeast Transformation II kits. NcAAs were purchased from the indicated companies: *p*-acetyl-L-phenylalanine (SynChem), *p*-azido-L-phenylalanine (Chem-Impex International), *O*-methyl-L-tyrosine (Chem-Impex International), *p*-propargyloxy-L-phenylalanine (Iris Biotech), 4-azidomethyl-L-phenylalanine (SynChem), 4-iodo-L-phenylalanine (AstaTech), 3,4-dihydroxy-L-phenylalanine (Alfa Aesar), 4-borono-L-phenylalanine (Acros Organics), 3-Amino-L-tyrosine (Bachem), 4-Amino-L-phenylalanine (Bachem), (*S*)-2-amino-6-((2-azidoethoxy)carbonylamino)hexanoic acid (Iris Biotech), (*S*)-2-amino-6-(((prop-2-yn-1-yloxy)carbonyl)amino)hexanoic acid (AstaTech), *N^ε^*-Boc-L-lysine (Chem-Impex International), *N^ε^*-azido-L-lysine (Chem-Impex International), *N^ε^,N^ε^* -dimethyl-L-lysine (Chem-Impex International), and L-α-aminocaprylic acid (Acros Organics).

### Media preparation and yeast strain construction

The preparation of liquid and solid media was performed as described previously.^*31*^ Unless otherwise noted, all SD-SCAA and SG-SCAA media used here were prepared without tryptophan (TRP), leucine (LEU) or uracil (URA). The strain RJY100 was constructed using standard homologous recombination approaches as described previously.^*41*^

### Preparing ncAA acid liquid stocks

All ncAA stocks were prepared at a final concentration of 50 mM L-isomer. DI water was added to the solid ncAA to approximately 90% of the final volume needed to make the stock, and 6.0 N NaOH was used as needed to fully dissolve the ncAA powder in the water. Water was added to the final volume and the solution was sterile filtered through a 0.2 micron filter. OmeY was pH adjusted to 7 prior to sterile filtering. No pH adjustment of additional ncAAs was performed unless otherwise noted. Filtered solutions were stored at 4 °C for up to four weeks for less labile ncAAs; for more labile ncAAs (AzF, BPhe, DOPA), 50 mM stocks were made immediately prior to induction.

### Reporter and secretion plasmid construction

The pCTCON2-FAPB2.3.6 and pCTCON2-FAPB2.3.6L1TAG reporter constructs have been previously described.^*31*^ pCTCON2-FAPB2.3.6 was used as a WT control to compare TAG-containing samples against, and was not used for library construction or sorting. The construction of reporter constructs pCHA-Donkey1.1-TAA and pCHA-Donkey1.1-H54TAG-TAA has also been reported previously.^*63*^

### EcTyrRS and EcLeuRS Library construction and characterization

Construction of plasmids pRS315-KanRmod-AcFRS^*63*^ and pRS315-EcLeuRS^*31*^ has been described previously. The EcLeuRS gene and cognate tRNA_CUA_, as well as the constitutive promoters for each gene, were PCR-amplified from pRS315-EcLeuRS and cloned into pRS315-KanRmod-AcFRS digested with SacI and PstI via Gibson assembly. The resulting plasmid was sequence verified and named pRS315-KanRmod-EcLeuRS.

Primers containing degenerate codons were used to amplify the aaRS genes from parent plasmids pRS315-KanRmod-EcLeuRS (EcLeuRS library) or pRS315-KanRmod-AcFRS (EcTyrRS library). Seven positions in the EcTyrRS active site were chosen for mutation: Y37, L71, Q179, N182, F183, L186, and Q195 (SI Tables 1 and 2). A separate primer with only the WT tyrosine codon at position Y37 was also used. The AcFRS gene contained a preexisting D165G mutation. An additional mutation, I7M, was inadvertently introduced when a primer containing that mutation was received and used for PCR amplification of the gene. However, a side-by-side comparison of AcFRS with and without the I7M mutations showed that the activity of AcFRS was not significantly affected by the presence of the mutation (SI Fig. 9). pRS315-KanRmod-AcFRS was digested with restriction enzymes NcoI and NdeI and the PCR-amplified EcTyrRS gene fragments were concentrated using Pellet Paint® NF Co-precipitant according to the manufacturer’s protocols. Similarly, seven positions in the EcLeuRS active site were chosen for mutation: M40, L41, T252, S496, Y499, Y527, and H537, with only an alanine mutation at position T252 (SI Tables 3, 4, and 7). pRS315-KanRmod-EcLeuRS was digested with restriction enzymes NcoI and NdeI and the PCR-amplified EcLeuRS gene fragments were concentrated using Pellet Paint® NF Co-precipitant according to the manufacturer’s protocols. A control DNA sample for each aaRS library that contained only doubly digested vector and no insert DNA was prepared similarly.

Electrocompetent *S. cerevisiae* RJY100 were prepared as described previously and electroporation protocols for transforming the library DNA were also followed as previously described.^*64*^ Electroporated cells were plated to determine transformation efficiency on SD-SCAA (−TRP −LEU −URA),^*64*^ and the remainder of cells were recovered in 100 mL SD-SCAA (−TRP −LEU −URA) supplemented with penicillin-streptomycin. One electroporation was performed for each library (EcTyrRS, EcLeuRS Library A, and EcLeuRS Library B). Dilutions plated on solid media to evaluate transformation efficiency were grown at 30 °C for 3–4 days and colonies were counted from multiple dilutions to approximate the number of individual transformants.^*64*^ The remainder of the libraries were grown at 30 °C with shaking at 300 rpm to saturation, then expanded into 1 L SD-SCAA (−TRP −LEU −URA) and grown at 30 °C overnight. The 1 L cultures were centrifuged at 3,214 rcf for 30 min and the supernatant was decanted. The cell pellets were resuspended in 60% glycerol to a final concentration of 15% glycerol, aliquoted to cryogenic vials, and stored at −80 °C. At least 5 × 10^9^ cells were stored per vial. A portion of each library was passaged separately in 5 mL SD-SCAA (−TRP −LEU −URA) cultures to characterize the naïve libraries using flow cytometry and to isolate and sequence the aaRS plasmids from several individual colonies (see “AaRS characterization post-FACS” for details).

### Error-prone PCR (epPCR) library construction and characterization

EcTyrRS mutant A-DOPARS-8 was used as a template for epPCR. epPCR was performed by combining 5 μL 10X ThermoPol Buffer, 1 μL 10 mM dNTPs, 5 μL 20 μM or 100 μM dPTP, 5 μL 20 μM or 100 μM 8-oxo-dGTP, 1 μL Taq polymerase, 1 μL DNA template (1 ng total), and 2.5 μL of each forward and reverse primer at 10 μM to amplify across the entire aaRS gene, as well as 27 μL sterile water to bring the total volume to 50 μL. Two concentrations (20 μM or 100 μM) of mutagenic dNTPs were used to vary the number of mutations made across the aaRS. Reactions were run on the thermal cycler at 95 °C for 500 s followed by 16 cycles of 95 °C for 45 s, 60 °C for 30 s, 72 °C for 135 s. Once cycles were complete, samples underwent a 10 min 72 °C final extension and hold at 4 °C until they were removed from the thermal cycler.

Following PCR with mutagenic dNTPs, each gene was amplified again via PCR at a higher volume to prepare enough DNA for electroporation into yeast. PCR was performed by combining 20 μL 10X ThermoPol Buffer, 4 μL 10 mM dNTP, 4 μL Taq polymerase, 10 μL epPCR-mutated DNA template, and 2 μL of each forward and reverse primer at 100 μM to amplify across the entire aaRS gene, as well as 158 μL sterile water to bring the total volume to 200 μL. Reactions were run on the thermal cycler at 95 °C for 180 s followed by 30 cycles of 95 °C for 45 s, 55 °C for 30 s, 72 °C for 135 s. Once cycles were completed, samples underwent a 10 min 72 °C final extension and hold at 4 °C until they were removed from the thermal cycler.

Digested pRS315-KanRmod-AcFRS vectors (still containing tRNA_CUA_^Tyr^) were prepared in the same manner as for the EcTyrRS saturation mutagenesis library (see above). For each of the DOPARS epPCR libraries, the following masses of DNA were concentrated using the Pellet Paint procedure: 4 μg epPCR-amplified aaRS, 1 μg double digested pRS315-KanRmod vector, and 1 μg pCTCON2-FAPB2.3.6L1TAG. Preparation of electrocompetent cells, electroporations, and subsequent characterization proceeded in the same manner as construction of EcTyrRS and EcLeuRS libraries (see above).

### Yeast transformations, propagation, and induction

Reporter plasmids pCTCON2-FAPB2.3.6L1TAG or pCTCON2-FAPB2.3.6 (TRP marker) and pRS315 plasmids from which the aaRS/tRNA pairs are expressed (LEU marker) were cotransformed into Zymo competent RJY100 cells, plated on solid SD-SCAA media (−TRP −LEU −URA), and grown at 30°C until colonies appeared (3 days). WT controls containing only pCTCON2-FAPB2.3.6 were transformed similarly into Zymo competent RJY100 cells, plated on solid SD-SCAA media (−TRP −URA), and grown at 30°C until colonies appeared (3 days). Inoculation and propagation in liquid SD media and induction in SG media have been described in detail elsewhere.^*31*^ Briefly: Three separate transformant colonies (biological triplicates) were inoculated from each plate except the WT control, where only one colony was inoculated, in 5 mL SD media of the same composition as the plates from transformations. All liquid cultures were supplemented with penicillin-streptomycin to prevent bacterial contamination. Liquid cultures were grown to saturation and then diluted to OD_600_ = 1 in 5 mL of the same media. The diluted cultures were grown to OD_600_ 2–5 (4–6 h at 30 °C with shaking) and then induced in 2 mL SG media at OD_600_ 1. Induction cultures with no ncAA and 1 mM final concentration of each respective ncAA were prepared for each replicate. The WT control was only induced with no ncAA. In the case of the epPCR aaRSs, induction cultures with 0.1 mM final concentration of the respective ncAA were also prepared. Induced cultures were incubated at 20 °C with shaking at 300 rpm for 16 h.

For library propagation and induction, the steps were identical but in 100 mL media inoculated from a 2 mL glycerol stock of the library with propagation and inductions in 100 mL in order to preserve the full diversity of the library.

### Flow cytometry data collection and analysis

Freshly induced samples were labeled in 1.7 mL microcentrifuge tubes or 96-well V-bottom plates. Flow cytometry was performed on an Attune NxT flow cytometer (Life Technologies) at the Tufts University Science and Technology Center. Detailed protocols describing the antibody labeling process have been described elsewhere.^*31*^ Primary and secondary antibody names and dilutions can be found in SI Table 13. Flow cytometry data analysis was performed using FlowJo and Microsoft Excel. For a sample gating strategy for flow cytometry data analysis in FlowJo, see SI Figure 10. Detailed descriptions of the calculations for RRE and MMF with corresponding error propagation have been described previously.^*31, 32*^

### Fluorescence-activated cell sorting

Aminoacyl-tRNA synthetase library populations were induced and labeled using identical methods to the “Flow cytometry data collection and analysis” section.^*31*^ For naïve library screens, 6 × 10^7^ cells were used for antibody labeling and antibody/PBSA volumes for primary and secondary labeling were adjusted accordingly. A number of cells was used that was at minimum ten times larger than the library population being sorted for all subsequent screens. Cell pellets were resuspended in ice-cold PBSA immediately prior to sorting. Samples were sorted using a FACSAria™ III (Becton, Dickinson and Company) flow cytometer at the Tufts University Flow Cytometry Core. Some first round sorts of Pooled Library A were performed on a MoFlo (Beckman-Coulter) and sorted in all subsequent rounds on the FACSAria™ III. Sorted samples were collected in 14 mL culture tubes containing 1 mL SD-SCAA (−TRP −LEU −URA) supplemented with penicillin-streptomycin. Following sorting, the sides of the culture tubes were washed with an additional 1 mL SD-SCAA (−TRP −LEU −URA), then transported back to the main laboratory facilities. An additional 3 mL SD-SCAA (−TRP −LEU −URA) was added to each sample and cultures were then grown at 30 °C with shaking at 300 rpm until saturated (2–3 days). Subsequent flow cytometry characterization was performed on each sorted population before the following round of screening (see above for details).

### AaRS characterization post-FACS

Once library populations exhibiting low stop codon readthrough in the presence of only cAAs and higher stop codon readthrough in the presence of ncAA(s) were isolated, aaRS plasmid DNA was purified using a Zymoprep Yeast Plasmid Miniprep II kit with slightly modified manufacturer’s protocols. 500 μL of 5 mL cultures library populations that had been previously propagated for flow cytometry characterization were diluted into 4.5 mL of the same media, supplemented with penicillin-streptomycin. Cultures were grown for 4 h at 30 °C with shaking at 300 rpm. 1 mL of each culture was removed to a microcentrifuge tube and pelleted at 13,000 rcf for 30 s. Supernatant was aspirated and each pellet was resuspended in 200 μL Solution I with 6 μL reconstituted zymolase from the Zymo kit. Each sample was vortexed briefly and then incubated at 37 °C with shaking at 300 rpm overnight or up to 24 h. 200 μL of Solution II from the Zymo kit was added and tubes were inverted to mix. 400 μL of Solution III was added and tubes were inverted to mix. Samples were pelleted at 13,000 rcf for 30 min to separate out cell debris. The supernatant was transferred to Epoch Life Science *E. coli* DNA purification columns and purified using the manufacturer’s protocols. DNA was eluted in 40 μL sterile water and then transformed into chemically competent *E. coli* DH5alphaZ1 cells and plated on LB media with 34 μg/mL kanamycin. 10-12 individual colonies were inoculated into separate 5 mL LB cultures with 50 μg/mL kanamycin and grown overnight at 37 °C with shaking at 300 rpm. Cultures were miniprepped using an Epoch *E. coli* GenCatch™ Plasmid DNA Mini-Prep Kit and submitted for sequencing.

### Secreted protein expression

Secretion plasmids pCHA-Donkey1.1-TAA and pCHA-Donkey1.1-L93TAG-TAA (constructed and described previously by Van Deventer and coworkers^*63*^) were transformed into chemically competent RJY100 using same method as described in the “Yeast transformations, propagation, and induction” section. Plasmid pCHA-Donkey1.1-TAA was not transformed with a second plasmid, while pCHA-Donkey1.1-L93TAG-TAA was co-transformed with pRS315 plasmids expressing the aaRS/tRNA_CUA_ pairs to be evaluated. All cells transformed with the WT construct were grown on or in SD-CAA during propagation and induced in YPG. All other cells cotransformed with pCHA constructs were grown on or in SD-SCAA (−TRP −LEU −URA) for growth and induced in YPG. All liquid cultures for growth and induction were supplemented with penicillin-streptomycin. Transformations, selection, and initial propagation in liquid media were essentially identical to pCTCON2 transformations in the “Yeast transformations, propagation, and induction” section. Following the initial inoculation of a single transformant colony into 5 mL SD selective media, cultures were grown at 30 °C for 24–48 h with shaking at 300 rpm. Each 5 mL culture was diluted into 45 mL SD media and grown for 24 h at 30 °C with shaking at 300 rpm. 50 mL cultures were then diluted into 450 mL SD media and grown for 24 h at 30 °C with shaking at 300 rpm. For inductions, 500 mL cultures were transferred to Nalgene centrifuge bottles and pelleted for 10 min at 2400 rcf. Supernatant was decanted and pellets were resuspended in 1 L YPG media with 1 mM final concentration of the corresponding ncAA and induced for 96 h at 20 °C with shaking at 300 rpm.

### Secreted protein purification

Induced cultures were centrifuged in Nalgene centrifuge bottles at 3,214 rcf for 30 min and the supernatant was sterile filtered along with a volume of 10X PBS buffer (1.37 M NaCl, 27 mM KCl, 100 mM Na_2_HPO_4_, and 18 mM KH2PO4 at pH 7.4) to bring the filtrate to a final concentration of 1X PBS. Protein purification columns were washed with 25 mL 1X PBS pH 7.0 before adding 2 mL Protein A resin (GenScript) and then washed with 25 mL 1X PBS. The equilibrated supernatant was added to each column and passed over the resin twice to improve protein capture efficiency. The resin was then washed three times with 1X PBS. Proteins were eluted in 7 mL 100 mM glycine pH 3.0 into a conical tube containing 0.7 mL 1M Tris pH 8.5 for immediate neutralization. Elution fractions were buffer exchanged into sterile water chilled to 4 °C using 15 mL 30 kDA molecular weight cutoff devices (Millipore Sigma). Buffer-exchanged proteins were flash frozen in 50% v/v glycerol and stored at −80 °C.

### Tryptic digests and sample preparation for mass spectrometry

Frozen glycerol stocks of secreted proteins (Donkey1.1) were thawed in a water bath at room temperature and buffer exchanged into sterile water chilled to 4 °C using 0.5 mL 30 kDA molecular weight cutoff devices (Millipore Sigma). A volume of protein corresponding to 10 μg total Donkey1.1 was boiled at 100 °C for 5 min and then cooled to room temperature on the benchtop. 1 μg Trypsin Gold (Promega) was added and samples were incubated at 37 °C overnight. Following tryptic digests, all samples were desalted using ZipTip C18 (Millipore Sigma) protocols, flash frozen in liquid nitrogen, and then shipped on dry ice to a mass spectrometry core facility. Trypsinized proteins were not resolved via chromatography prior to mass spectrometry analysis.

## Acknowledgments

This work was supported in part by a grant from the Army Research Office (W911NF-16-1-0175), in part by a grant from the National Institute of General Medical Sciences of the National Institutes of Health (1R35GM133471), and in part by Tufts University startup funds (to J.A.V.). FACS performed at the Tufts Laser Cytometry core facility was partly funded by the National Institutes of Health (1S10OD016196-01). J.T.S. was supported in part by an NSF Graduate Research Fellowship (ID: 2016231237). The content is solely the responsibility of the authors and does not necessarily represent the official views of the Army Research Office, the National Institutes of Health, the National Science Foundation, or Tufts University. We would like to thank Stephen Kwok and Allen Parmelee for their expertise and assistance with FACS and Michael Ream for his help with error-prone PCR experiments. Additionally, we would like to thank Dr. Richard Cook, Heather Amoroso, and Alla Leshinsky at the Koch Institute Biopolymers and Proteomics Core for their assistance with mass spectrometry data collection. Finally, we thank Dr. Rafael Alcala-Toraño and Rebecca Hershman for their invaluable editing contributions and feedback.

## Author Contributions

Conceptualization, J.T.S. and J.A.V.; Methodology, J.T.S. and J.A.V.; Investigation, J.T.S.; Writing – Original Draft, J.T.S.; Writing – Review & Editing, J.T.S. and J.A.V.; Funding Acquisition, J.A.V.; Supervision, J.A.V.

## Data Availability Statement

The flow cytometry and sequencing datasets generated during and/or analyzed during the current study are available from the corresponding author on reasonable request.

## Availability of Materials

All materials from this study are available from the corresponding author upon reasonable request.

