## Supplementary Information for "High-throughput aminoacyl-tRNA synthetase engineering for genetic code expansion in yeast"

**Supplementary discussion**

1. RRE and MMF calculations
2. MALDI mass spectrometry analysis

**Supplementary figures**

1. Depiction of which saturation mutagenesis aaRS libraries were used for different types of sorts
2. Comparison of the activity of two borono-phenylalynyl-tRNA synthetases (BPheRSs)
3. Flow cytometry plots of Track 2 polyspecific clones PolyT2RS-5, PolyT2RS-7, and A-APheRS-4
4. MALDI mass spectrometry characterization of ncAA-containing tryptic digested peptide fragments.
5. Performance of aaRS mutants isolated using random mutagenesis
6. Error for RRE and MMF values in Figure 6 of the main text
7. Flow cytometry characterization between FACS rounds for polyspecificity Track 2
8. A Track 1 polyspecific sort population induced in the presence of 21 ncAAs
9. RRE and MMF for TyrAcFRS with and without the I7M mutation
10. Gating strategy for FlowJo analysis of flow cytometry data

**Supplementary tables**

1. EcTyrRS saturation mutagenesis library design
2. EcTyrRS saturation mutagenesis library primers
3. EcLeuRS saturation mutagenesis library design
4. EcLeuRS Library A primers
5. EcTyrRS library sequence characterization
6. EcLeuRS Library A sequence characterization
7. EcLeuRS Library B primers
8. EcLeuRS Library B sequence characterization
9. Sequences of all EcTyrRS variants isolated via FACS for individual ncAA, specificity, and polyspecificity sorts
10. Sequences of all EcLeuRS variants isolated via FACS for individual ncAA, specificity, and polyspecificity sorts.
11. Expected peptide sizes for the tryptic digest fragment containing the H54TAG codon from the scFv-Fc form of Donkey 1.1
12. Sequences of DOPARS clone variants isolated following error-prone mutagenesis and FACS
13. Flow cytometry primary and secondary labeling conditions and reagents

### Supplementary Discussion

Relative readthrough efficiency (RRE) is a measure of C-terminal detection divided by N-terminal detection for a protein containing a UAG codon over the same terminus detections for the equivalent wild-type (WT) protein (Equation 1). Maximum misincorporation frequency (MMF) is the RRE evaluated in the absence of ncAAs during induction over the RRE evaluated in the presence of ncAAs during induction (Equation 2). RRE and MMF can be used with any protein that has dual-terminus detection, whether the detected entities are fluorescent proteins, epitope tags, or alternative methods that allow for fluorescent labels to be attached and detected on a spectrophotometric plate reader or flow cytometer. It is not possible to perform traditional statistical analyses on calculated RRE and MMF values, as all the data from biological triplicates is aggregated during the calculations; separate RRE and MMF values could be defined for individual sets of samples, but the pairings of WT and UAG samples would be arbitrary. Statistical analyses could be performed on the averaged MFI values of the detection of the C-terminus of the reporter proteins, but these values are not normalized to the total amount of expressed reporter or to the WT protein expression, eliminating important controls to ensure that readthrough efficiency, not changes in gene expression, are being evaluated.[1]

Eqn. 1 $$RRE = \frac{UAG\ C\ terminus\ detection}{UAG\ N\ terminus\ detection} \bigg/ \frac{WT\ C\ terminus\ detection}{WT\ N\ terminus\ detection}$$

Eqn. 2 $$MMF = \frac{RRE_{-ncAA}}{RRE_{+ncAA}}$$

Incorporation of ncAAs was confirmed via MALDI mass spectrometry of an enzymatically digested reporter protein (SI Fig. 4). Notably, low yields due to inefficient translation with some combinations of aaRSs and ncAAs imposed limits on the resolution of mass spectrometry data achievable in some samples. Expected peptide sizes and peptide sizes that could appear due to cAA misincorporation can be found in SI Table 11. Expected and actual masses are reported directly on the MS spectra. Peptide masses at 2210.1, 2282.2, and 2298.2 Da are due to trypsin autolysis.[2] The expected peptide masses of interest appeared in most samples, though some peptides showed clear signs of ncAA degradation, which has been noted in previous literature. For example, both A-DOPARS-4 and DOPARS-0.1-10 have a low peak at approximately 2310 Da, which we attribute to a dehydration event that results in removal of one of the hydroxyl groups (SI Fig. 4D and E). Similarly, BPhe can be dehydrated or doubly dehydrated to 2322 Da and 2304 Da, respectively.[3, 4] In this case, the peptide masses corresponding to intact BPhe, the single dehydration product, and the double dehydration product were observed at 2337.9, 2322.2, and 2303.8 Da, respectively (SI Fig. 4F). Additional peptide masses appeared in the A-ATyrRS-1 sample at 2334 and 2347 Da that we attribute to peptide masses for missed trypsin cleavage sites (SI Fig. 4G). The peptide corresponding to the 2334 Da peak is VSNKALPAPIEKTISKAKGQPR and the peptide corresponding to the 2347 Da peak is EYKCKVSNKALPAPIEKTISK. The peptide peak at 2333 Da that appears in the spectrum for B-LysAlkRS-3 is a known peptide mass that appears when tryptophan is encoded at the H54TAG position by the OTS (SI Fig. 4J). For B-BocKRS-2 induced in the presence of 1 mM BocK, the peak at 2381 Da is also indicative of a peptide mass that arises from a single missed cleavage site: ASQSISSYLNWYQQKPGKAPK (SI Fig. 4K). Similar to the B-LysAlkRS-3 sample, the peak at approximately 2333 Da is likely due to misincorporation of tryptophan by B-BockRS-2. The PolyT2RS-5 sample induced in the presence of 1 mM AzF also showed a peak at 2309.3 Da indicative of degradation of AzF to APhe (expected peptide mass of 2309.2 Da, SI Fig. 4O).[5-7] The expected peptide peak corresponding to intact AzF also appeared at 2335.2 Da. The PolyT2RS-5 sample induced in the presence of 1 mM AzMF showed the expected peptide peak at 2349.3 Da, with additional degradation peaks observed from the degradation of the azide group (SI Fig. 4R).

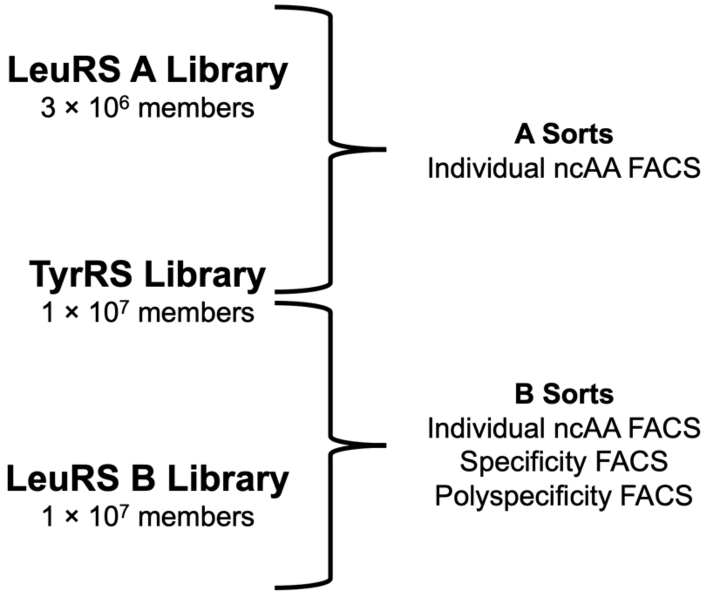


Supplementary Figure 1. Depiction of which saturation mutagenesis aaRS libraries were used for different types of sorts. Pooled TyrRS and LeuRS Library A were used solely for sorting for activity with individual ncAAs. Pooled TyrRS and LeuRS Library B were used for sorts with individual ncAAs as well as sorts for desirable specificity profiles (i.e. specificity and polyspecificity sorts).

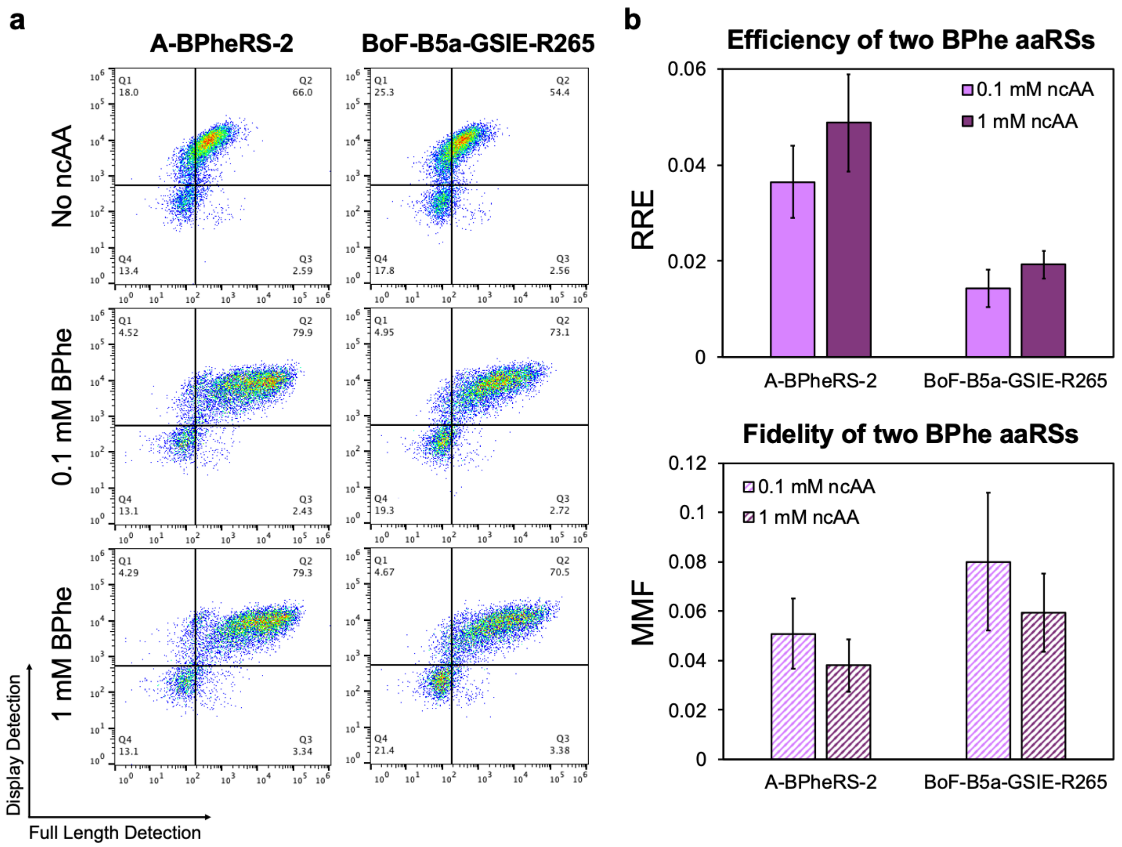


**c**

| Clone | Mutations |
|---|---|
| A-BPheRS-2 | Y37G, L71T, Q179N, D182S, F183V, L186E, Q195I |
| BoF-B5a-GSIE-R265 | Y37G, D182S, F183I, L186E, D265R |

Supplementary Figure 2. Comparison of the activity of two borono-phenylalynyl-tRNA synthetases (BPheRSs). **a**, Flow cytometry dot plots of two BPheRSs: A-BPheRS-2 from pooled TyrRS and LeuRS Library A screens and BoF-B5a-GSIE-R265 from Chatterjee and coworkers.[3] BPheRSs were induced in the absence of ncAAs and in the presence of 0.1 mM and 1 mM BPhe. Display (N-terminal) detection is on the Y axis and full-length (C-terminus) detection is on the X axis. **b**, Quantitative evaluation of the efficiency (relative readthrough efficiency, RRE) and fidelity (maximum misincorporation frequency, MMF) of the two BPheRSs at two concentrations of BPhe. **c**, Mutations relative to WT TyrRS for each of the two BPheRSs.

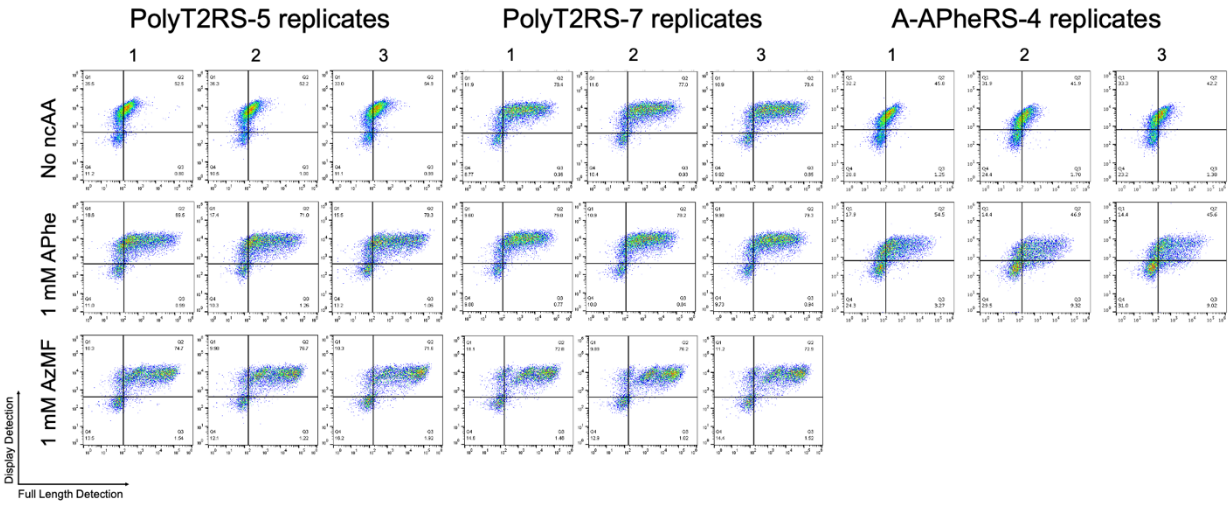


Supplementary Figure 3. Flow cytometry plots of Track 2 polyspecific clones PolyT2RS-5, PolyT2RS-7, and A-APheRS-4. Both polyspecificity clones were induced in the absence of ncAAs and in the presence of 1 mM APhe and 1 mM AzMF. A-APheRS-4 was only induced in the absence of ncAAs and in the presence of 1 mM APhe.

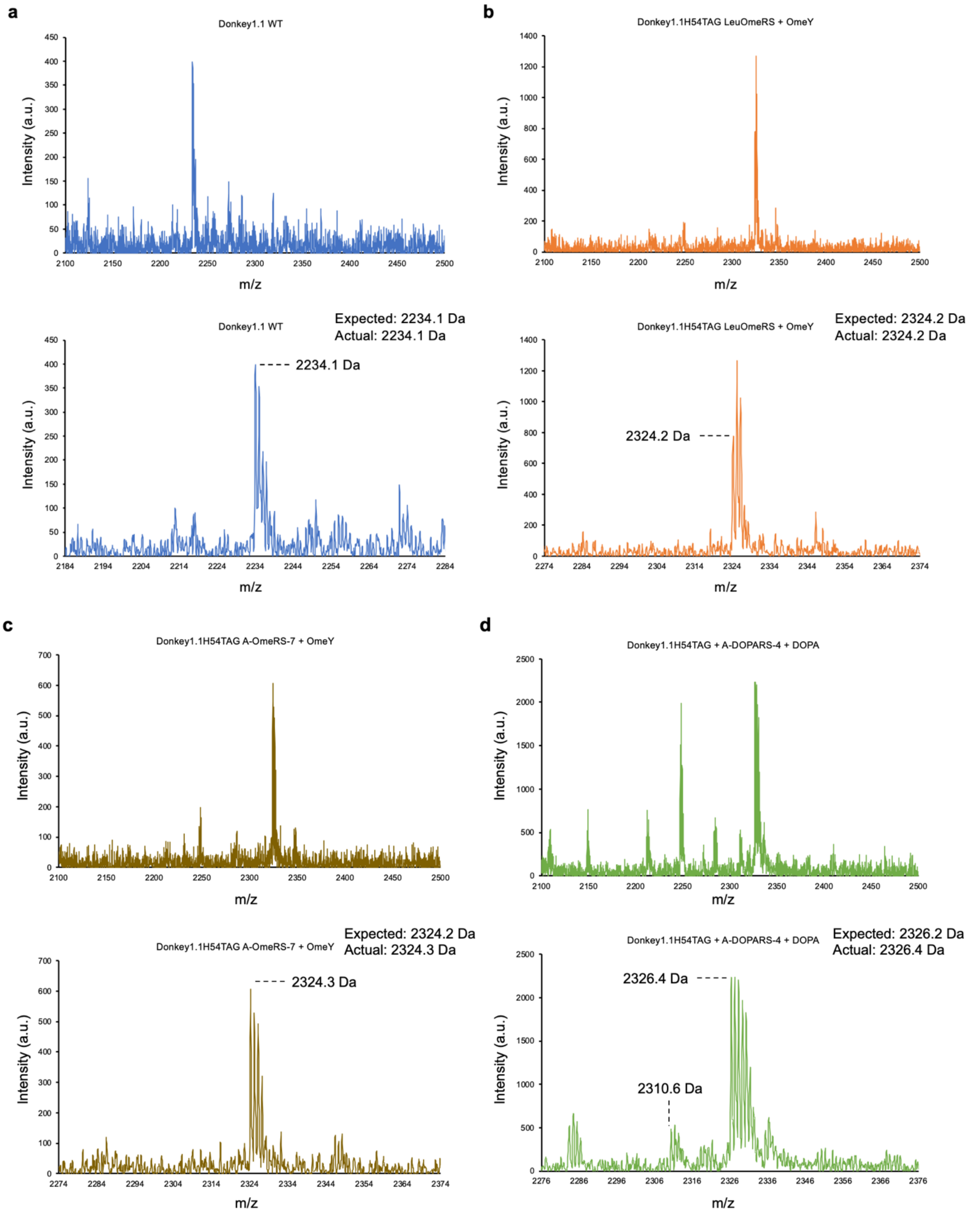

a
Donkey1.1 WT
Intensity (a.u.)
m/z
Donkey1.1 WT
Expected: 2234.1 Da
Actual: 2234.1 Da
2234.1 Da
b
Donkey1.1H54TAG LeuOmeRS + OmeY
Donkey1.1H54TAG LeuOmeRS + OmeY
Expected: 2324.2 Da
Actual: 2324.2 Da
2324.2 Da
c
Donkey1.1H54TAG A-OmeRS-7 + OmeY
Donkey1.1H54TAG A-OmeRS-7 + OmeY
Expected: 2324.2 Da
Actual: 2324.3 Da
2324.3 Da
d
Donkey1.1H54TAG + A-DOPARS-4 + DOPA
Donkey1.1H54TAG + A-DOPARS-4 + DOPA
Expected: 2326.2 Da
Actual: 2326.4 Da
2326.4 Da
2310.6 Da

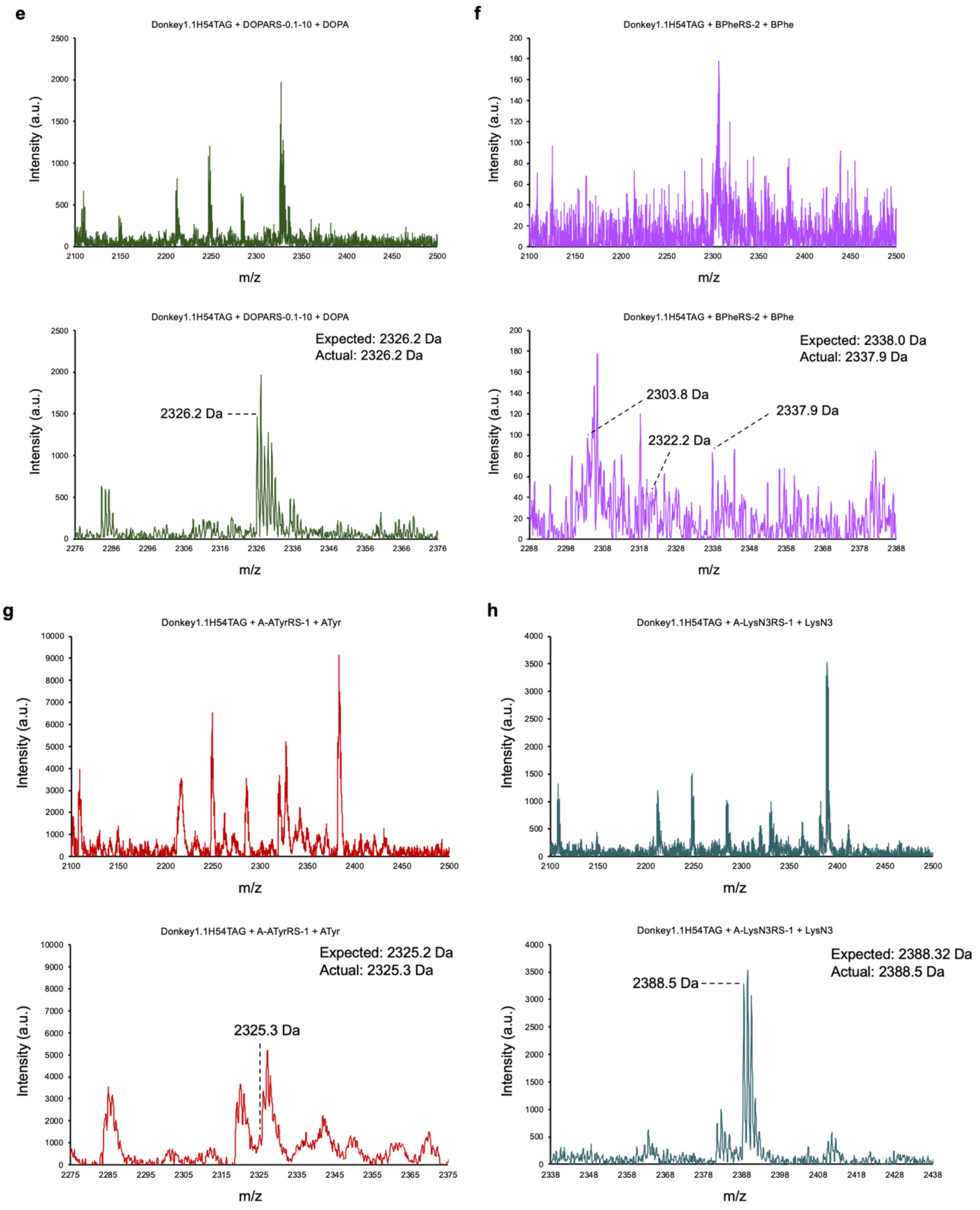
e
Donkey1.1H54TAG + DOPARS-0.1-10 + DOPA
Intensity (a.u.)
m/z
Donkey1.1H54TAG + DOPARS-0.1-10 + DOPA
Expected: 2326.2 Da
Actual: 2326.2 Da
2326.2 Da
Intensity (a.u.)
m/z
f
Donkey1.1H54TAG + BPheRS-2 + BPhe
Intensity (a.u.)
m/z
Donkey1.1H54TAG + BPheRS-2 + BPhe
Expected: 2338.0 Da
Actual: 2337.9 Da
2303.8 Da
2337.9 Da
2322.2 Da
Intensity (a.u.)
m/z
g
Donkey1.1H54TAG + A-ATyrRS-1 + ATyr
Intensity (a.u.)
m/z
Donkey1.1H54TAG + A-ATyrRS-1 + ATyr
Expected: 2325.2 Da
Actual: 2325.3 Da
2325.3 Da
Intensity (a.u.)
m/z
h
Donkey1.1H54TAG + A-LysN3RS-1 + LysN3
Intensity (a.u.)
m/z
Donkey1.1H54TAG + A-LysN3RS-1 + LysN3
Expected: 2388.32 Da
Actual: 2388.5 Da
2388.5 Da
Intensity (a.u.)
m/z

**i**

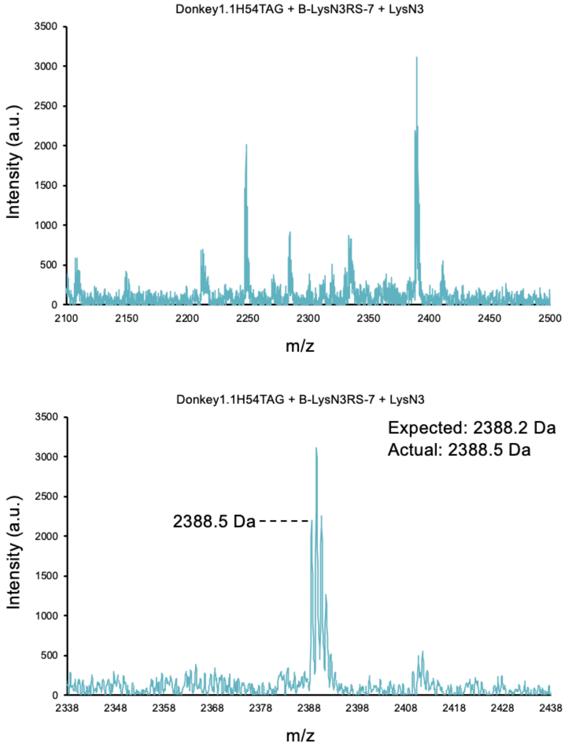
Donkey1.1H54TAG + B-LysN3RS-7 + LysN3
Intensity (a.u.)
m/z
Donkey1.1H54TAG + B-LysN3RS-7 + LysN3
Expected: 2388.2 Da
Actual: 2388.5 Da
2388.5 Da
Intensity (a.u.)
m/z

**j**

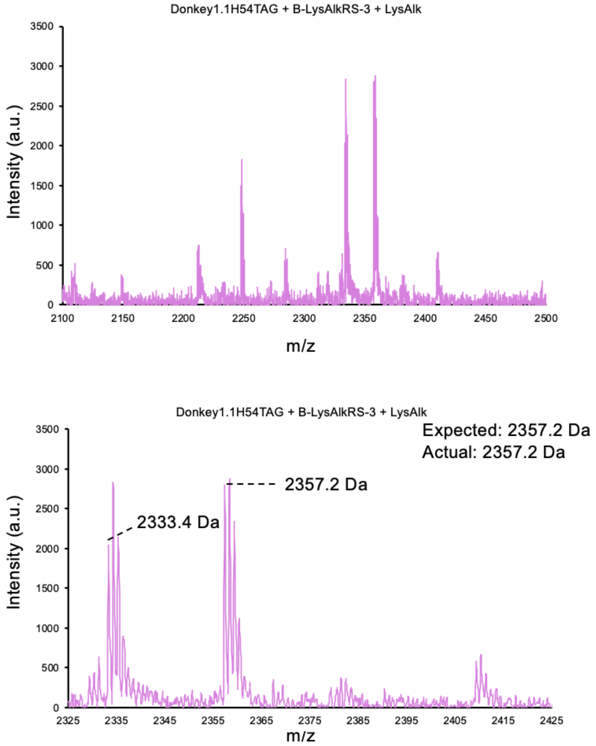
Donkey1.1H54TAG + B-LysAlkRS-3 + LysAlk
Intensity (a.u.)
m/z
Donkey1.1H54TAG + B-LysAlkRS-3 + LysAlk
Expected: 2357.2 Da
Actual: 2357.2 Da
2357.2 Da
2333.4 Da
Intensity (a.u.)
m/z

**k**

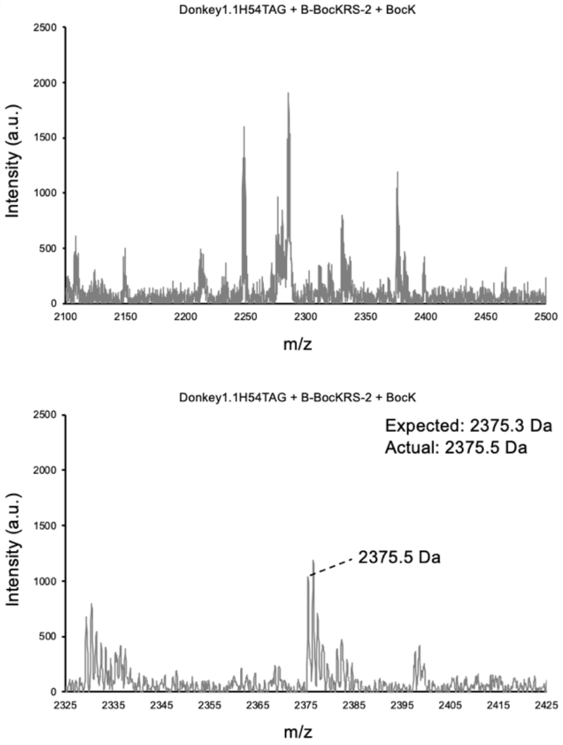
Donkey1.1H54TAG + B-BocKRS-2 + BocK
Intensity (a.u.)
m/z
Donkey1.1H54TAG + B-BocKRS-2 + BocK
Expected: 2375.3 Da
Actual: 2375.5 Da
2375.5 Da
Intensity (a.u.)
m/z

**l**

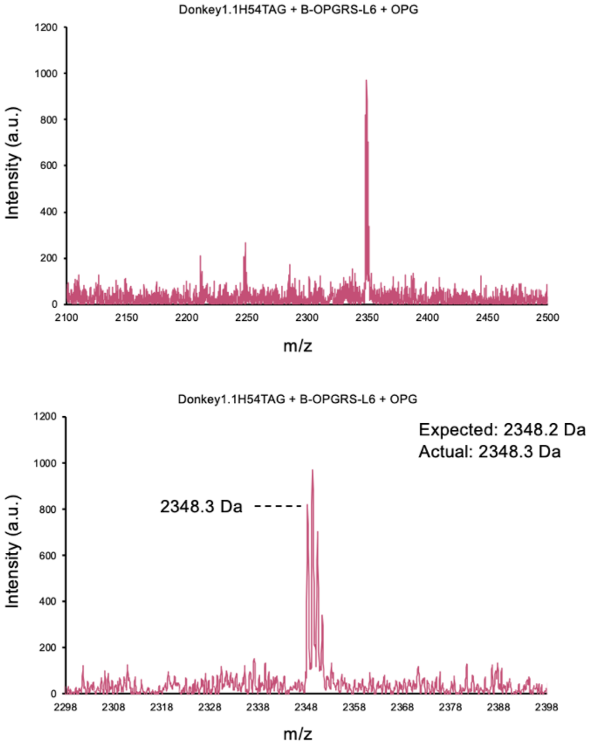
Donkey1.1H54TAG + B-OPGRS-L6 + OPG
Intensity (a.u.)
m/z
Donkey1.1H54TAG + B-OPGRS-L6 + OPG
Expected: 2348.2 Da
Actual: 2348.3 Da
2348.3 Da
Intensity (a.u.)
m/z

m

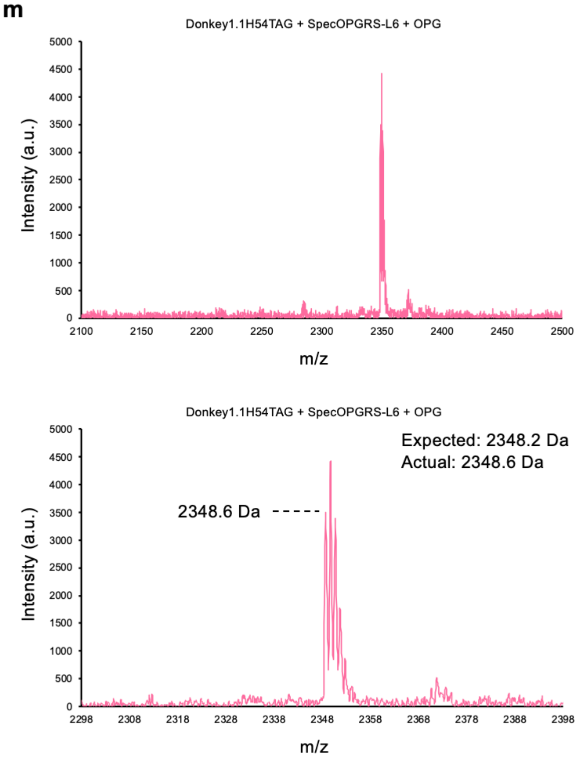
Donkey1.1H54TAG + SpecOPGRS-L6 + OPG
Intensity (a.u.)
m/z
Donkey1.1H54TAG + SpecOPGRS-L6 + OPG
Expected: 2348.2 Da
Actual: 2348.6 Da
2348.6 Da
Intensity (a.u.)
m/z

n

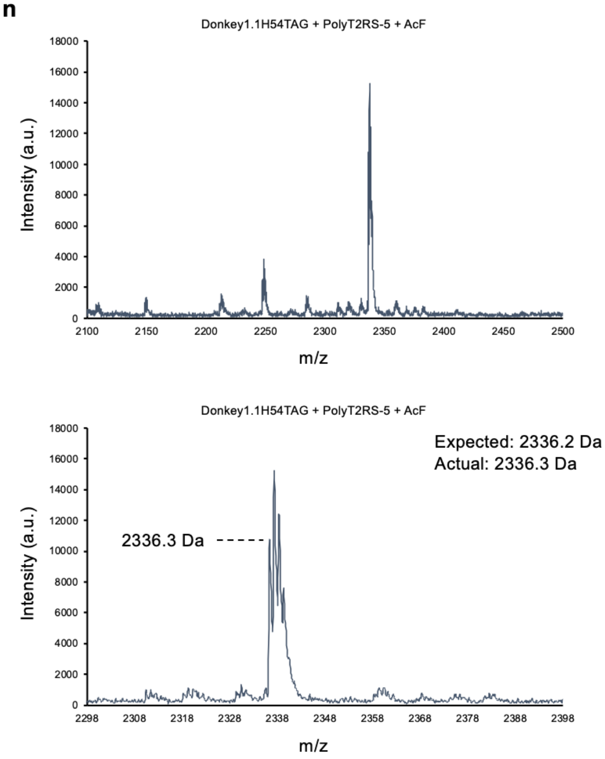
Donkey1.1H54TAG + PolyT2RS-5 + AcF
Intensity (a.u.)
m/z
Donkey1.1H54TAG + PolyT2RS-5 + AcF
Expected: 2336.2 Da
Actual: 2336.3 Da
2336.3 Da
Intensity (a.u.)
m/z

o

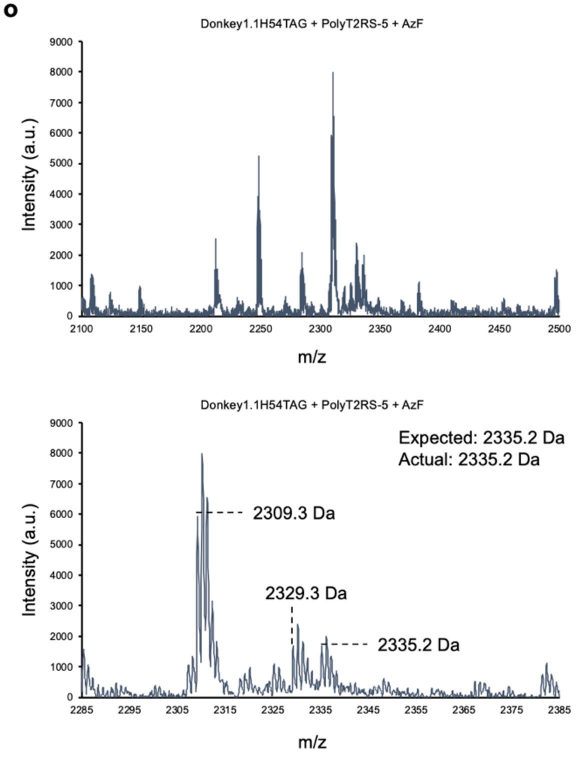
Donkey1.1H54TAG + PolyT2RS-5 + AzF
Intensity (a.u.)
m/z
Donkey1.1H54TAG + PolyT2RS-5 + AzF
Expected: 2335.2 Da
Actual: 2335.2 Da
2309.3 Da
2329.3 Da
2335.2 Da
Intensity (a.u.)
m/z

p

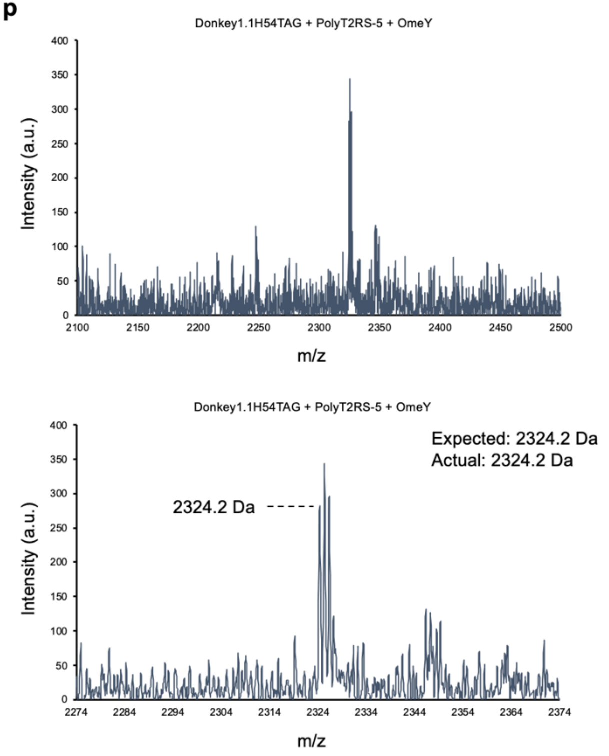
Donkey1.1H54TAG + PolyT2RS-5 + OmeY
Intensity (a.u.)
m/z
Donkey1.1H54TAG + PolyT2RS-5 + OmeY
Expected: 2324.2 Da
Actual: 2324.2 Da
2324.2 Da
Intensity (a.u.)
m/z

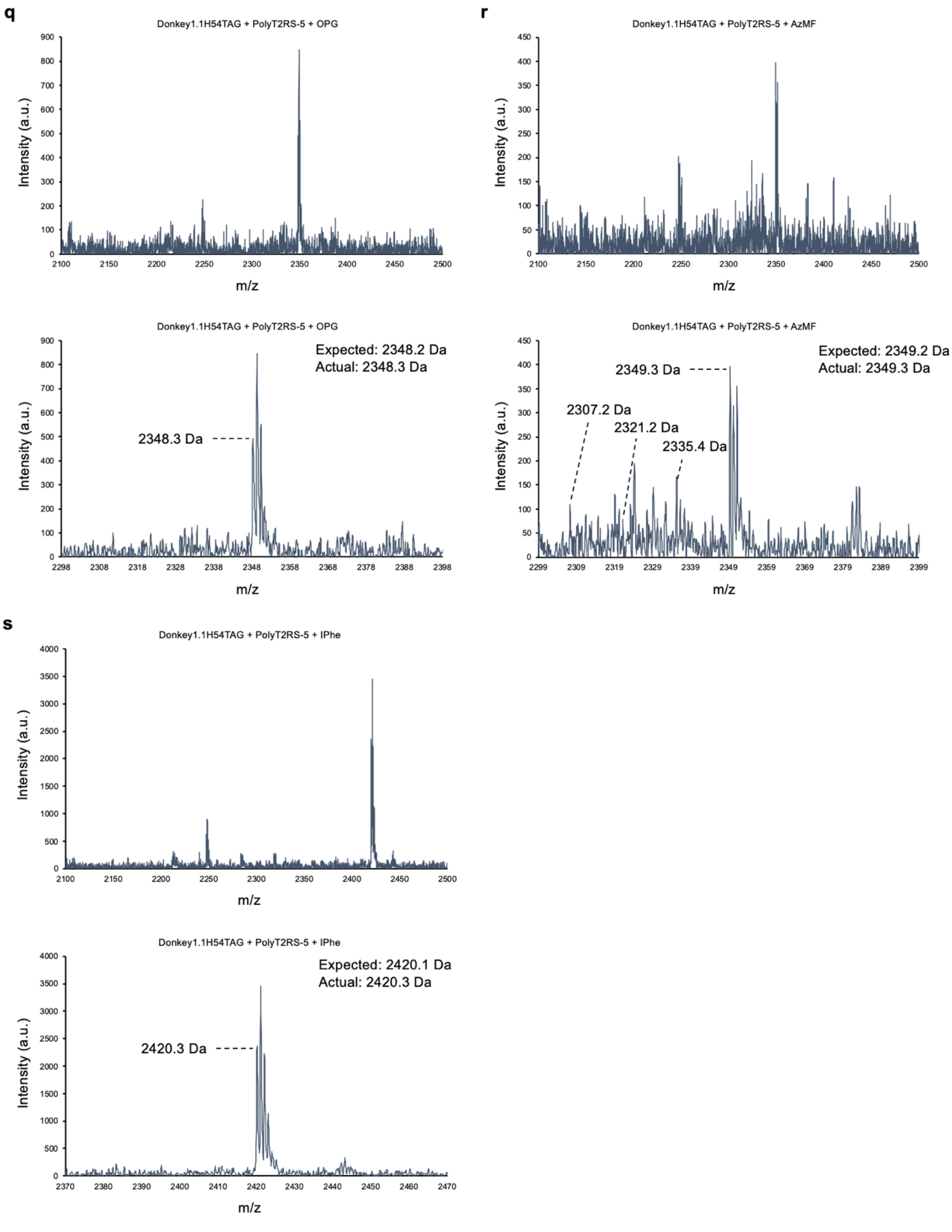


Supplementary Figure 4. MALDI mass spectrometry characterization of ncAA-containing tryptic digested peptide fragments. **a**, Wildtype (WT) Donkey1.1 reporter protein used to evaluate ncAA incorporation via MALDI MS. **b**, Donkey1.1-H54TAG reporter protein with LeuOmeRS induced with 1 mM OmeY. **c**, Donkey1.1-H54TAG with A-OmeRS-7 induced with 1 mM OmeY. **d**, Donkey1.1-H54TAG with A-DOPARS-4 induced with 1 mM DOPA. **e**, Donkey1.1-H54TAG with DOPARS-0.1-10 induced with 1 mM DOPA. **f**, Donkey1.1-H54TAG with A-BPheRS-2 induced with 1 mM BPhe. **g**, Donkey1.1-H54TAG with A-ATyrRS-1 induced with 1 mM ATyr. **h**, Donkey1.1-H54TAG with A-LysN3RS-1 induced with 1 mM LysN3. **i**, Donkey1.1-H54TAG with B-LysN3RS-7 induced with 1 mM LysN3. **j**, Donkey1.1-H54TAG with B-LysAlkRS-3 induced with 1 mM LysAlk. **k**, Donkey1.1-H54TAG with B-BocKRS-2 induced with 1 mM BocK. **l**, Donkey1.1-H54TAG with B-OPGRS-L6 induced with 1 mM OPG. **m**, Donkey1.1-H54TAG with SpecOPGRS-3 induced with 1 mM OPG. **n**, Donkey1.1-H54TAG with PolyT2RS-5 induced with 1 mM AcF. **o**, Donkey1.1-H54TAG with PolyT2RS-5 induced with 1 mM AzF. **p**, Donkey1.1-H54TAG with PolyT2RS-5 induced with 1 mM OmeY. **q**, Donkey1.1-H54TAG with PolyT2RS-5 induced with 1 mM OPG. **r**, Donkey1.1-H54TAG with PolyT2RS-5 induced with 1 mM AzMF. **s**, Donkey1.1-H54TAG with PolyT2RS-5 induced with 1 mM IPhe.

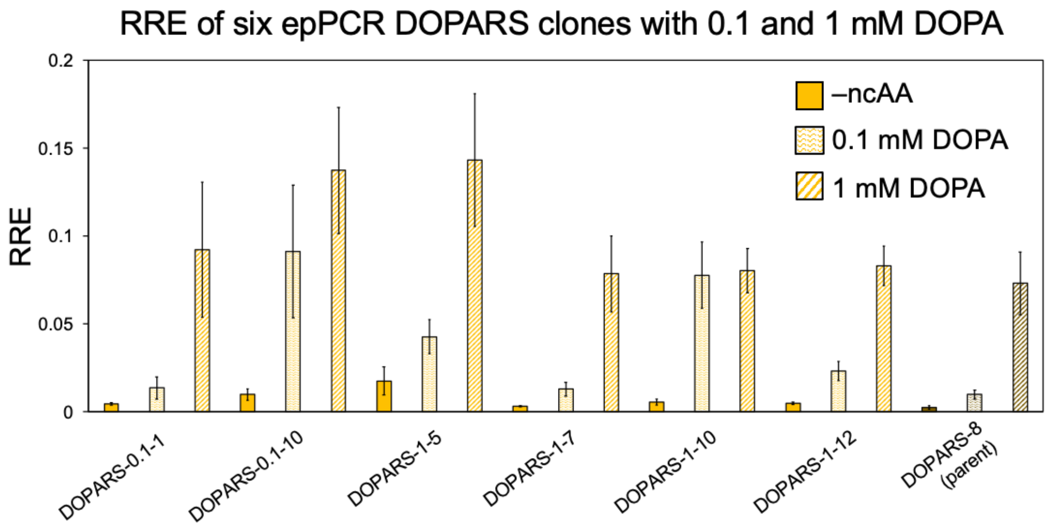


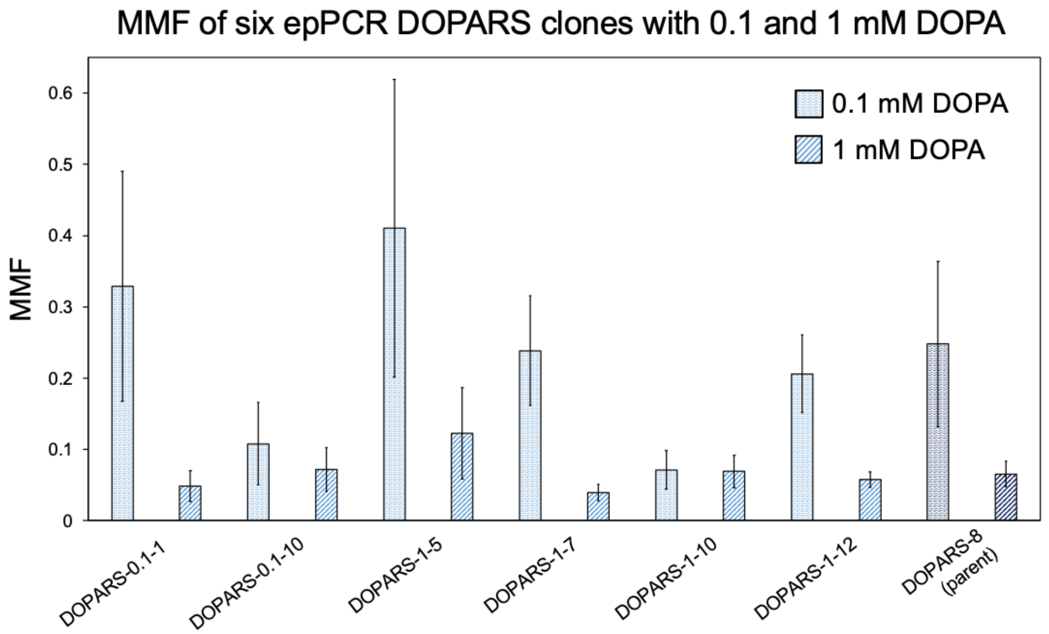


Supplementary Figure 5. Performance of aaRS mutants isolated using random mutagenesis. Top: RRE of six epPCR mutants that outperform the parent DOPARS at 0.1 mM, 1 mM, or both concentrations of DOPA. Bottom: MMF of six epPCR mutants that are comparable to the parent DOPARS fidelity of DOPA incorporation at both 0.1 and 1 mM DOPA. All RRE and MMF values were derived from samples in biological triplicate and error bars represent the standard deviation of triplicate values that was propagated during RRE and MMF calculations.

**a**

**RRE of aaRSs with aromatic ncAAs error**

| | –ncAA | AcF | AzF | OmeY | OPG | AzMF | IPhe | DOPA | BPhe | ATyr | APhe |
|---|---|---|---|---|---|---|---|---|---|---|---|
| **A-OmeRS-7** | 0.00014 | 0.080 | 0.27 | 0.12 | 0.056 | 0.00044 | 0.026 | 0.0012 | 0.00057 | 0.00099 | 0.00113 |
| **DOPARS-0.1-10** | 0.00058 | 0.0024 | 0.0016 | 0.0017 | 0.0013 | 0.00041 | 0.00034 | 0.050 | 0.0023 | 0.00103 | 0.00097 |
| **A-BPheRS-2** | 0.00034 | 0.00064 | 0.00056 | 0.00068 | 0.00057 | 0.00029 | 0.00016 | 0.00076 | 0.048 | 0.00045 | 0.00072 |
| **B-OPGRS-L6** | 0.00032 | 0.039 | 0.040 | 0.079 | 0.052 | 0.016 | 0.096 | 0.0018 | 0.00223 | 0.00184 | 0.00435 |
| **SpecOPGRS-3** | 0.0010 | 0.0015 | 0.0017 | 0.0012 | 0.052 | 0.00060 | 0.00059 | 0.0015 | 0.00146 | 0.00093 | 0.00085 |
| **PolyT2RS-5** | 0.00015 | 0.048 | 0.063 | 0.052 | 0.054 | 0.011 | 0.094 | 0.00043 | 0.00034 | 0.0003 | 0.00321 |
| **PolyT2RS-7** | 0.0026 | 0.028 | 0.017 | 0.18 | 0.041 | 0.0080 | 0.051 | 0.0041 | 0.00364 | 0.00459 | 0.00317 |
| **B-LysAlkRS-3** | 0.0041 | 0.0090 | 0.013 | 0.011 | 0.011 | 0.016 | 0.068 | 0.0072 | 0.01127 | 0.00736 | 0.00407 |
| **B-BocKRS-2** | 0.00041 | 0.054 | 0.090 | 0.13 | 0.062 | 0.024 | 0.15 | 0.0036 | 0.00036 | 0.00041 | 0.00043 |

**MMF of aaRSs with aromatic ncAAs error**

| | AcF | AzF | OmeY | OPG | AzMF | IPhe | DOPA | BPhe | ATyr | APhe |
|---|---|---|---|---|---|---|---|---|---|---|
| **A-OmeRS-7** | 0.00064 | 0.0022 | 0.0013 | 0.0040 | 0.095 | 0.0081 | 0.087 | 0.060 | 0.17 | 0.14 |
| **DOPARS-0.1-10** | 0.089 | 0.16 | 0.14 | 0.21 | 0.30 | 0.51 | 0.0049 | 0.22 | 0.13 | 0.25 |
| **A-BPheRS-2** | 0.15 | 0.11 | 0.16 | 0.29 | 0.35 | 0.33 | 0.24 | 0.0040 | 0.21 | 0.27 |
| **B-OPGRS-L6** | 0.0026 | 0.0027 | 0.0043 | 0.0029 | 0.030 | 0.0023 | 0.21 | 0.32 | 0.43 | 0.11 |
| **SpecOPGRS-3** | 0.34 | 0.33 | 0.46 | 0.016 | 0.79 | 0.65 | 0.30 | 0.42 | 0.45 | 0.55 |
| **PolyT2RS-5** | 0.0027 | 0.0031 | 0.0027 | 0.0015 | 0.011 | 0.0022 | 0.10 | 0.13 | 0.15 | 0.062 |
| **PolyT2RS-7** | 0.014 | 0.048 | 0.021 | 0.020 | 0.071 | 0.0071 | 0.15 | 0.27 | 0.24 | 0.36 |
| **B-LysAlkRS-3** | 0.15 | 0.096 | 0.067 | 0.075 | 0.050 | 0.038 | 0.10 | 0.13 | 0.14 | 0.13 |
| **B-BocKRS-2** | 0.0015 | 0.0020 | 0.0028 | 0.0016 | 0.015 | 0.0026 | 0.092 | 0.36 | 0.29 | 0.35 |

**b**

**RRE of aaRSs with aliphatic ncAAs error**

| | –ncAA | LysN3 | LysAlk | BocK | AzK | DMK | AC |
|---|---|---|---|---|---|---|---|
| **PolyT2RS-5** | 0.00026 | 0.00027 | 0.00021 | 0.00030 | 0.00035 | 0.00029 | 0.00027 |
| **PolyT2RS-7** | 0.0032 | 0.0075 | 0.0021 | 0.0024 | 0.0049 | 0.0042 | 0.025 |
| **B-LysAlkRS-3** | 0.0041 | 0.020 | 0.024 | 0.011 | 0.0021 | 0.0035 | 0.0083 |
| **B-BocKRS-2** | 0.00042 | 0.0057 | 0.0011 | 0.033 | 0.00038 | 0.0013 | 0.0043 |
| **B-LysN3RS-7** | 0.0035 | 0.051 | 0.0022 | 0.0020 | 0.0013 | 0.0021 | 0.0033 |
| **LeuOmeRS** | 0.00027 | 0.00022 | 0.00018 | 0.00016 | 0.00026 | 0.00023 | 0.00036 |
| **A-OmeRS-7** | 0.00016 | 0.00034 | 0.00034 | 0.0003 | 0.00031 | 0.00031 | 0.00025 |
| **B-OPGRS-L6** | 0.00018 | 0.00060 | 0.00046 | 0.00047 | 0.00014 | 0.00039 | 0.00022 |

**MMF of aaRSs with aliphatic ncAAs error**

| | LysN3 | LysAlk | BocK | AzK | DMK | AC |
|---|---|---|---|---|---|---|
| **PolyT2RS-5** | 0.24 | 0.29 | 0.51 | 0.23 | 0.27 | 0.37 |
| **PolyT2RS-7** | 0.12 | 0.35 | 0.49 | 0.24 | 0.36 | 0.039 |
| **B-LysAlkRS-3** | 0.011 | 0.018 | 0.068 | 0.11 | 0.10 | 0.053 |
| **B-BocKRS-2** | 0.0091 | 0.018 | 0.0030 | 0.078 | 0.046 | 0.012 |
| **B-LysN3RS-7** | 0.013 | 0.35 | 0.35 | 0.31 | 0.33 | 0.28 |
| **LeuOmeRS** | 0.16 | 0.34 | 0.46 | 0.23 | 0.28 | 0.22 |
| **A-OmeRS-7** | 0.074 | 0.086 | 0.10 | 0.12 | 0.16 | 0.13 |
| **B-OPGRS-L6** | 0.092 | 0.087 | 0.13 | 0.11 | 0.17 | 0.15 |

Supplementary Figure 6. Error for RRE and MMF values in Figure 6 of the main text. Comprehensive evaluation of aaRS activity with aromatic and aliphatic ncAAs. **a**, RRE and MMF error values for nine aaRSs with 10 aromatic ncAAs. **b**, RRE and MMF error values for eight aaRSs with six aliphatic ncAAs. All RRE and MMF values were derived from samples evaluated in biological triplicate with error bars representing the standard deviation of triplicate values that was propagated during RRE and MMF calculations. Error was calculated from measurements performed in biological triplicate as described in *Materials and Methods*.

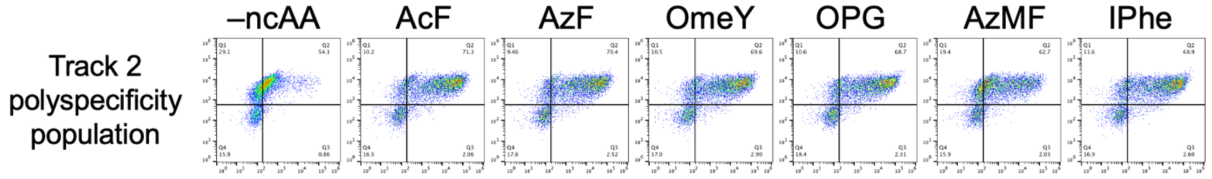


Supplementary Figure 7. Flow cytometry characterization between FACS rounds for polyspecificity Track 2. The pooled naïve EcTyRS and B EcLeuRS libraries underwent the following sorts: 1 mM AzF (round 1), 1 mM AcF (round 2), negative (round 3) and 1 mM AzMF (round 4) prior to flow cytometry characterization in this figure. The population was induced in the absence of ncAAs and in the presence of 1 mM AcF, AzF, OmeY, OPG, AzMF, and IPhe.

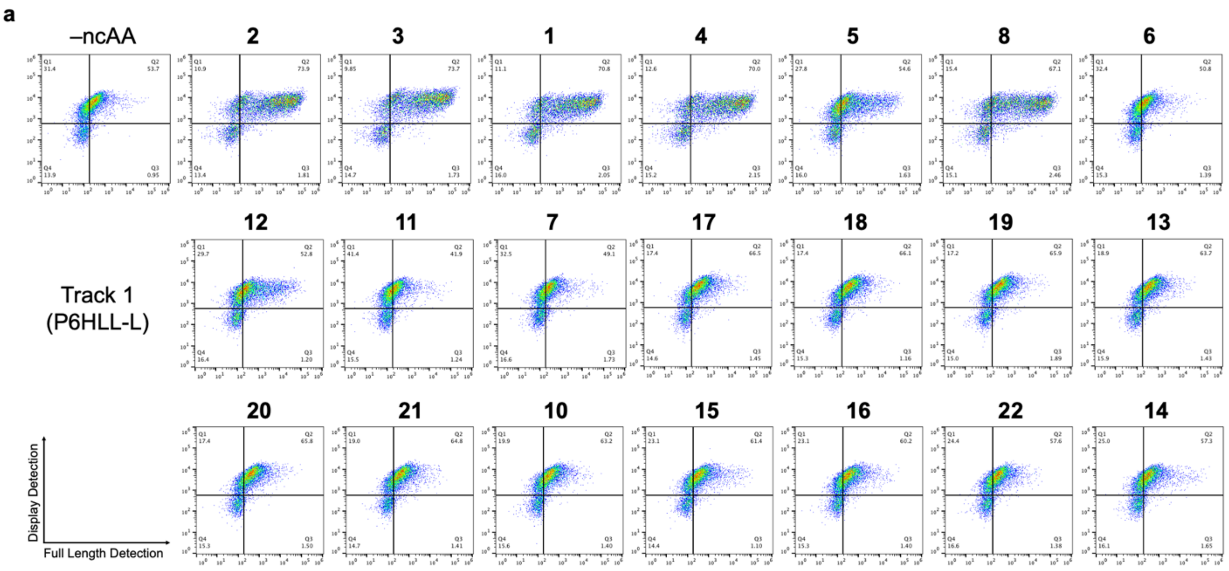


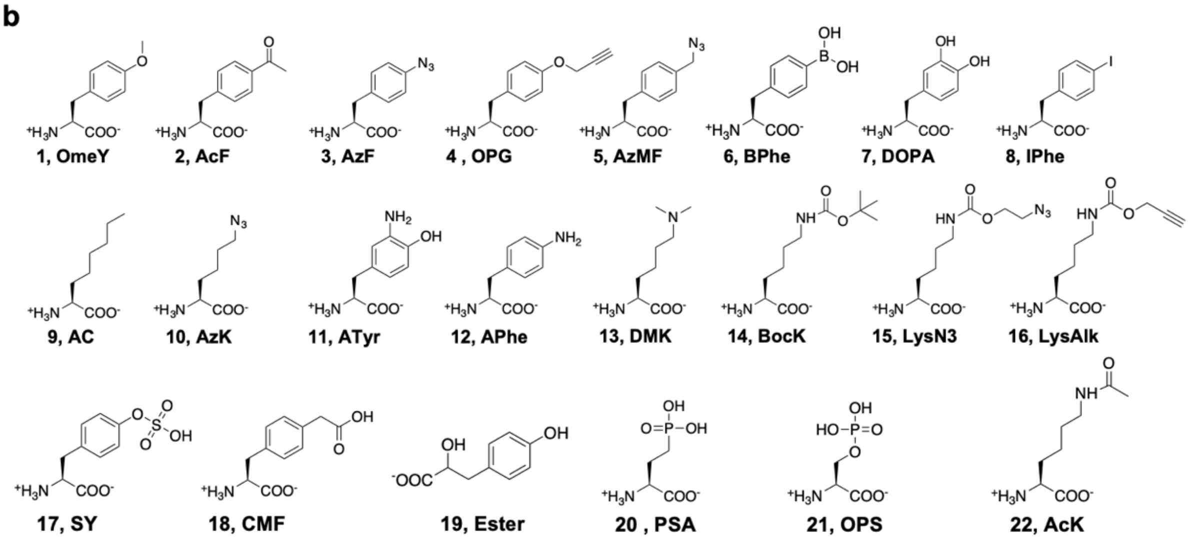


Supplementary Figure 8. A Track 1 polyspecific sort population induced in the presence of 21 ncAAs. **a**, Flow cytometry dot plots for a Track 1 polyspecificity sort (denoted as sort “P6HLL-L”) induced in the absence of ncAAs (–ncAA) and in the presence of 1 mM 21 ncAAs. **b**, Structures of ncAAs used in part a.

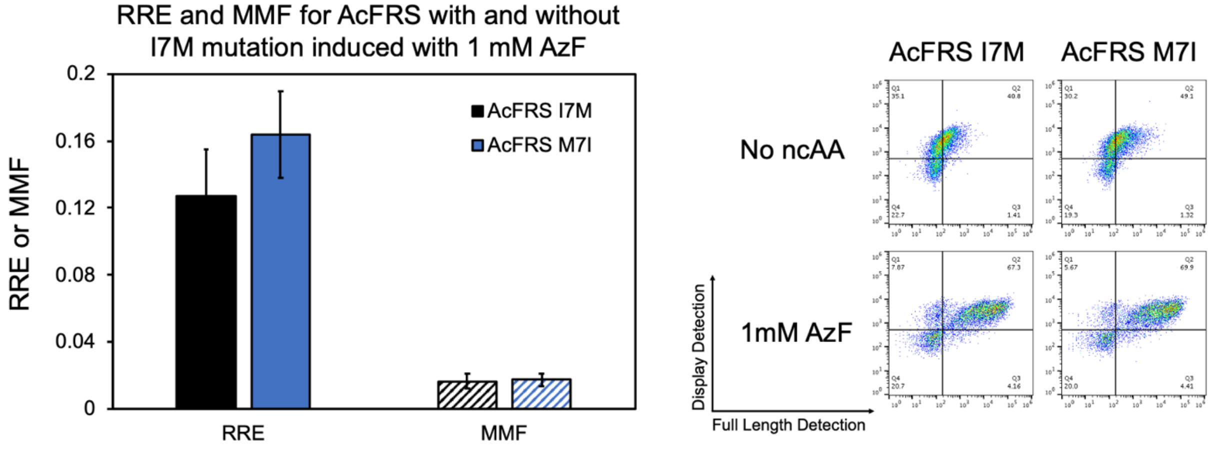


Supplementary Figure 9. RRE and MMF for TyrAcFRS with and without the I7M mutation. Qualitative flow cytometry plots are shown on the right. Samples were induced in the absence of ncAAs and in the presence of 1 mM AzF.

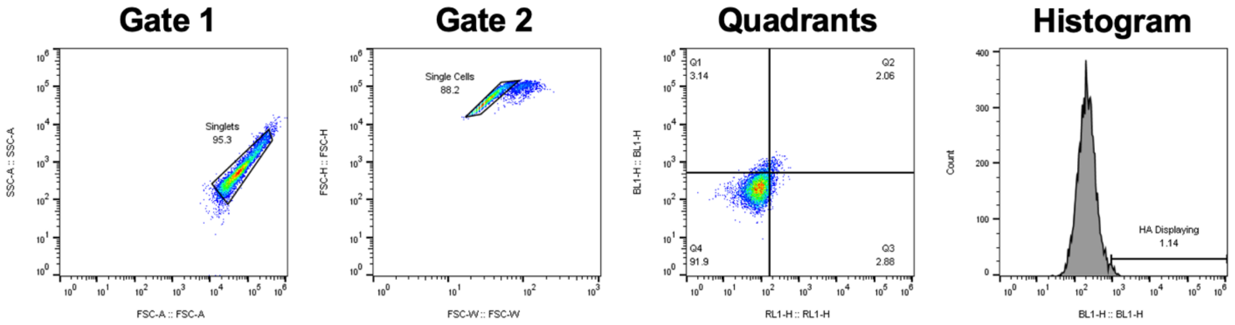


Supplementary Figure 10. Gating strategy for FlowJo analysis of flow cytometry data. For each individual day of a flow cytometry experiment, a new set of gates was applied to all samples. This was typically done by using a population of *S. cerevisiae* RJY100 that was unlabeled (i.e., no primary or secondary antibodies were used for this sample) to gate out live, single cells. For the first gate, SSC area versus FSC area, a gate was drawn around the population of cells that encompassed the living yeast cells (Gate 1). Gate 2 was drawn on the Gate 1 population on a FSC height versus FSC width plot to remove possible doublet, triplet, or larger groups of cells from the flow cytometry data analysis. Quadrants were placed on the Gate 2 cell population on a BL1 (blue laser, detection channel 1) height versus RL1 (red laser, detection channel 1) height plot. For all flow cytometry experiments, the blue laser was used to detect the N-terminal HA epitope tag and the red laser was used to detect the C-terminal c-Myc epitope tag. Additionally, for RRE and MMF data analysis, a gate was drawn on histogram of BL1 height. This gate was used to identify the population of cells exhibiting HA detection above autofluorescence levels.

Supplementary Table 1. *E. coli* tyrosyl-tRNA synthetase (EcTyrRS) saturation mutagenesis library design. The wild-type (WT) EcTyrRS residue and position are indicated that were mutated using degenerate codons VNK, RRT, or KYA. For position Y37, the WT codon TAT was also included. Degenerate codon VNK encodes the following residues: Leu, Pro, His, Gln, Arg, Ile, Met, Thr, Asn, Lys, Ser, Val, Ala, Asp, Glu, and Gly. Degenerate codon RRT encodes the following residues: Asn, Ser, Asp, and Gly. Degenerate codon KYA encodes the following residues: Leu, Ser, Val, and Ala. The numbers of possible codons at each position were used to determine the theoretical diversity for the EcTyrRS library.

| **Position** | **Codon** | **# Codons** |
|---|---|---|
| Y37 | VNK + TAT | 25 |
| L71 | VNK | 24 |
| Q179 | VNK | 24 |
| D182 | RRT | 4 |
| F183 | VNK | 24 |
| L186 | KYA | 4 |
| Q195 | VNK | 24 |
| **Theoretical Diversity** | | **1.3E8** |

Supplementary Table 2. EcTyrRS library primers used to mutate the residues indicated in SI Table 1. Degenerate codons are colored blue, and the WT codon at Y37 (TAT) is indicated in pink.

| Position | Forward | Reverse |
|---|---|---|
| Y37 | GAGCGACTGGCGCAAGGCCCGATCGCGCTC VNKTGCGGCTTCGATCCTACCGCTGACAGC | GCTGTCAGCGGTAGGATCGAAGCCGCAMNBG AGCGCGATCGGGCCTTGCGCCAGTCGCTC |
| Y37 | GAGCGACTGGCGCAAGGCCCGATCGCGCTC TATTGCGGCTTCGATCCTACCGCTGACAGC | GCTGTCAGCGGTAGGATCGAAGCCGCAATAG AGCGCGATCGGGCCTTGCGCCAGTCGCTC |
| L71 | CAGCAGGCGGGCCACAAGCCGGTTGCGVNK GTAGGCGGCGCGACGGGTCTGATTGGCGAC | GTCGCCAATCAGACCCGTCGCGCCGCCMNBT ACCGCAACCGGCTTGTGGCCCGCCTGCTG |
| Q179, D182, F183, L186 | CGTTCACTGAGTTTTCCTACAACCTGTTGVNK GGTTATRRTVNKGCCTGTKYAAACAAACAGTA CGGTGTGGTGCTGCAAATTG | CAATTTGCAGCACCACACCGTACTGTTTGTTT RMACAGGCMNBAYYATAACCMNBCAACAGGT TGTAGGAAAACTCAGTGAACG |
| Q195 | GTGCTAACAAACAGTACGGTGTGGTGCTGVN KATTGGTGGTTCTGACCAGTGGGGTAAC | GTTACCCCACTGGTCAGAACCACCAATMNBCA GCACCACACCGTACTGTTTGTTAGCAC |

Supplementary Table 3. *E. coli* leucyl-tRNA synthetase (EcLeuRS) saturation mutagenesis library design. The wild-type (WT) EcLeuRS residue and position are indicated that were mutated using degenerate codons VNK, RST, NNY, or NNT. For editing domain position T252, only a GCT mutation to alanine was allowed. Degenerate codon VNK encodes the following residues: Leu, Pro, His, Gln, Arg, Ile, Met, Thr, Asn, Lys, Ser, Val, Ala, Asp, Glu, and Gly. Degenerate codon RST encodes the following residues: Thr, Ala, Ser, and Gly. Degenerate codons NNY and NNT encode the following residues: Phe, Ser, Tyr, Cys, Leu, Pro, His, Arg, Ile, Thr, Asn, Val, Ala, Asp, and Gly. The numbers of possible codons at each position were used to determine the theoretical diversity for the EcLeuRS library.

| Position | Codon | # Codons |
|---|---|---|
| M40 | VNK | 24 |
| L41 | VNK | 24 |
| T252 | GCT | 1 |
| S496 | RST | 4 |
| Y499 | NNY | 32 |
| Y527 | NNT | 16 |
| H537 | NNT | 16 |
| **Theoretical Diversity** | | **1.9E7** |

Supplementary Table 4. EcLeuRS Library A primers. Original EcLeuRS library primers used to mutate the residues indicated in SI Table 3. Degenerate codons are colored blue, and the T252A mutation is colored orange. These primers were used to construct EcLeuRS Library A.

| Position | Forward | Reverse |
|---|---|---|
| M40, L41 | GCAAAGAGAAGTATTACTGCCTGTCTVNKVNKCCCTATCCTTCTGGTCGACTACACATG | CATGTGTAGTCGACCAGAAGGATAGGGMNBMNBAGACAGGCAGTAATACTTCTCTTTGC |
| T252 | CTGACCGTTTACACTACCCGCCCGGACGCTTTTATGGGTTGTACCTACCTGGCGGTAGC | GCTACCGCCAGGTAGGTACAACCCATAAAAGCGTCCGGGCGGGTAGTGTAAACGGTCAG |
| S496, Y499 | GAAACCGACACTTTCGACACCTTTATGGAGRSTTCCTGGNNYTATGCGCGCTACACTTGCCCGCAGTACAAAG | CTTTGTACTGCGGGCAAGTGTAGCGCGCATARNNCCAGGAASYCTCCATAAAGGTGTCGAAAGTGTCGGTTTC |
| Y527 | GCGGCTAACTACTGGCTGCCGGTGGATATCNNTATTGGTGGTATTGAACACGCCATTATG | CATAATGGCGTGTTCAATACCACCAATANNGATATCCACCGGCAGCCAGTAGTTAGCCGC |
| H537 | TATTGGTGGTATTGAACACGCCATTATGNNTCTGCTCTACTTCCGCTTCTTCCACAAAC | GTTTGTGGAAGAAGCGGAAGTAGAGCAGANNCATAATGGCGTGTTCAATACCACCAATA |

Supplementary Table 5. EcTyrRS library sequence characterization. 10 clones were chosen at random for Sanger sequence characterization after EcTyrRS library construction in yeast. All 10 clones were unique and had expected residues based on degenerate codon design at the seven positions mutated.

| WT Codon | Y37 | L71 | Q179 | D182 | F183 | L186 | Q195 |
|---|---|---|---|---|---|---|---|
| **Degenerate codon** | **VNK + TAT** | **VNK** | **GCT** | **RRT** | **VNK** | **KYA** | **VNK** |
| **1** | G | G | R | N | K | L | V |
| **2** | R | S | R | D | G | A | R |
| **3** | S | V | A | D | A | A | P |
| **4** | L | L | S | D | P | A | Q |
| **5** | D | M | E | D | R | A | L |
| **6** | M | V | L | N | I | L | V |
| **7** | H | D | T | G | M | A | P |
| **8** | S | M | R | S | H | L | P |
| **9** | G | V | N | S | D | A | V |
| **10** | N | V | R | D | N | A | I |

Supplementary Table 6. EcLeuRS Library A sequence characterization. 10 clones were chosen at random for Sanger sequence characterization after EcLeuRS library A construction in yeast. All 10 clones were unique but had predominantly WT residues at positions Y527 and H537, which was not expected based on the degenerate codon design.

| WT Codon | M40 | L41 | T252 | S496 | Y499 | Y527 | H537 |
|---|---|---|---|---|---|---|---|
| Degenerate codon | VNK | VNK | GCT | RST | NNY | NNT | NNT |
| 1 | N | T | A | T | T | Y | H |
| 2 | A | S | A | T | A | Y | H |
| 3 | V | E | A | T | A | Y | H |
| 4 | A | R | A | G | D | Y | F |
| 5 | G | A | A | T | P | S | A |
| 6 | N | R | A | A | A | Y | T |
| 7 | I | R | A | A | I | Y | L |
| 8 | G | H | A | A | H | V | S |
| 9 | N | R | A | T | S | V | Y |
| 10 | R | D | A | G | Y | Y | H |

Supplementary Table 7. EcLeuRS Library B primers. Modified EcLeuRS library primers used to mutate the residues indicated in SI Table 3. Degenerate codons are colored blue, and the T252A mutation is colored orange. These primers were used to construct EcLeuRS Library B.

| Position | Forward | Reverse |
|---|---|---|
| M40, L41 | GCAAAGAGAAGTATTACTGCCTGTCTVNKVNKCCCTATCCTTCTGGTCGACTACACATG | CATGTGTAGTCGACCAGAAGGATAGGGMNBMNBAGACAGGCAGTAATACTTCTCTTTGC |
| T252 | CTGACCGTTTACACTACCCGCCCGGACGCTTTTATGGGTTGTACCTACCTGGCGGTAGC | GCTACCGCCAGGTAGGTACAACCCATAAAAGCGTCCGGGCGGGTAGTGTAAACGGTCAG |
| S496, Y499 | GAAACCGACACTTTCGACACCTTTATGGAGRSTTCCTGGNNYTATGCGCGCTACACTTGCCCGCAGTACAAAG | CTTTGTACTGCGGGCAAGTGTAGCGCGCATARNNCCAGGAASYCTCCATAAAGGTGTCGAAAGTGTCGGTTTC |
| Y527, H537 | GCGGCTAACTACTGGCTGCCGGTGGATATCNNTATTGGTGGTATTGAACACGCCATTATGNNTCTGCTCTACTTCCGCTTCTTCCACAAACTG | CAGTTTGTGGAAGAAGCGGAAGTAGAGCAGANNCATAATGGCGTGTTCAATACCACCAATANNGATATCCACCGGCAGCCAGTAGTTAGCCGC |

Supplementary Table 8. EcLeuRS Library B sequence characterization. Nine clones were chosen at random for Sanger sequence characterization after EcLeuRS library B construction in yeast. Eight out of the nine clones were unique and had expected residues based on degenerate codon design at the seven positions mutated.

| WT Codon | M40 | L41 | T252 | S496 | Y499 | Y527 | H537 |
|---|---|---|---|---|---|---|---|
| Degenerate codon | VNK | VNK | GCT | RST | NNY | NNT | NNT |
| 1 | I | D | A | A | H | Y | S |
| 2 | G | S | A | G | I | H | F |
| 3 | G | H | A | S | D | S | C |
| 4 | A | T | A | T | Y | D | I |
| 5 | A | T | A | T | Y | D | I |
| 6 | L | N | A | A | P | T | N |
| 7 | R | V | A | S | A | V | S |
| 8 | D | H | A | A | S | A | H |
| 9 | M | L | A | G | N | S | S |

Supplementary Table 9. Sequences of all EcTyrRS variants isolated via FACS for individual ncAA, specificity, and polyspecificity sorts.

| TyrRS Position | Y37 | L71 | V72† | Q179 | D182 | F183 | L186 | Q195 |
|---|---|---|---|---|---|---|---|---|
| *Codon* | *VNK + TAT* | *VNK* | *N/A* | *VNK* | *RRT* | *VNK* | *KYA* | *VNK* |
| A-OmeRS-1 | L | V | V | Q | G | M | A | Q |
| A-OmeRS-2 | I | L | V | Q | G | M | A | Q |
| A-OmeRS-3 | G | R | V | P | S | P | L | S |
| A-OmeRS-4 | I | L | V | Q | G | M | A | Q |
| A-OmeRS-5 | I | L | V | Q | G | M | A | Q |
| A-OmeRS-6 | I | L | V | Q | G | M | A | Q |
| A-OmeRS-7 | L | L | V | Q | G | M | A | Q |
| A-OmeRS-8 | I | L | V | Q | G | M | A | Q |
| A-OmeRS-9 | I | L | V | Q | G | M | A | Q |
| A-OmeRS-10 | L | L | V | Q | G | M | A | Q |
| A-DOPARS-1 | E | L | V | Q | G | M | A | T |
| A-DOPARS-2 | L | L | V | G | D | T | V | L |
| A-DOPARS-3 | T | L | V | M | D | N | L | V |
| A-DOPARS-4 | Q | M | V | S | D | T | V | I |
| A-DOPARS-5 | E | V | V | Q | G | M | A | V |
| A-DOPARS-6 | E | L | V | Q | G | M | A | T |
| A-DOPARS-7 | T | I | V | M | D | M | A | K |
| A-DOPARS-8 | T | I | V | M | D | M | A | K |
| A-DOPARS-9 | E | V | M | Q | G | M | A | T |
| A-LysN3RS-1.1 | V | V | V | A | D | T | S | I |
| A-LysN3RS-1.2 | A | M | V | Q | D | T | V | V |
| A-LysN3RS-1.3 | L | V | V | N | D | I | L | Q |
| A-LysN3RS-1.5 | I | L | V | Q | D | G | L | Q |
| A-LysN3RS-1.9 | I | L | V | Q | D | V | A | L |
| A-LysN3RS-1.10 | A | M | V | Q | D | T | V | G |
| A-LysN3RS-1.11 | L | D | V | L | D | T | L | V |
| A-BPheRS-1 | E | I | V | L | S | I | S | E |
| A-BPheRS-2 | G | T | V | N | S | V | E | I |
| A-BPheRS-3 | A | L | V | P | D | T | L | A |
| A-BPheRS-4 | G | T | V | N | S | V | E | I |
| A-BPheRS-5 | M | V | R | Q | D | L | A | S |
| A-BPheRS-6 | V | V | L | V | D | T | A | L |
| A-BPheRS-7 | G | T | V | N | S | V | E | I |
| A-BPheRS-8 | L | V | V | N | D | L | S | Q |
| A-BPheRS-9 | G | T | V | N | S | V | E | I |
| A-BPheRS-10 | E | I | V | L | S | I | S | E |
| A-BPheRS-11 | G | T | V | N | S | V | E | I |
| A-BPheRS-12 | G | T | V | N | S | V | E | I |
| A-APheRS-1 | I | L | V | Q | G | M | A | Q |
| A-APheRS-2 | I | L | V | Q | G | M | A | Q |
| A-APheRS-3 | I | L | V | Q | G | M | A | Q |
| A-APheRS-4 | V | V | V | Q | G | M | A | Q |
| A-APheRS-5 | I | L | V | Q | G | M | A | Q |
| A-APheRS-6 | G | L | V | T | D | T | S | M |
| A-APheRS-7 | I | L | V | Q | G | M | A | Q |
| A-APheRS-8 | E | L | V | Q | G | M | A | S |
| A-APheRS-9 | I | L | V | Q | G | M | A | Q |
| A-APheRS-10 | T | V | S | N | D | I | L | G |
| A-APheRS-11 | I | L | V | Q | G | M | A | Q |
| A-APheRS-12 | I | L | V | Q | G | M | A | Q |
| A-ATyrRS-4 | L | L | V | M | D | Q | A | S |
| A-ATyrRS-6 | L | V | V | D | D | T | A | E |
| A-ATyrRS-7 | L | M | V | N | D | I | S | L |
| A-ATyrRS-8 | L | L | V | M | D | Q | A | S |
| A-ATyrRS-10 | E | V | V | H | D | V | L | I |

Supplementary Table 9 continued

| TyrRS Position | Y37 | L71 | V72† | Q179 | D182 | F183 | L186 | Q195 |
|---|---|---|---|---|---|---|---|---|
| *Codon* | *VNK + TAT* | *VNK* | *N/A* | *VNK* | *RRT* | *VNK* | *KYA* | *VNK* |
| B-OPGRS-H1 | V | V | T | Q | G | M | A | Q |
| B-OPGRS-H2 | I | L | V | Q | G | M | A | Q |
| B-OPGRS-L1 | I | L | V | Q | G | M | A | Q |
| B-OPGRS-L2 | I | L | V | Q | G | M | A | Q |
| B-OPGRS-L3 | I | L | V | Q | G | M | A | Q |
| B-OPGRS-L4 | I | L | V | Q | G | M | A | Q |
| B-OPGRS-L5 | I | L | V | Q | G | M | A | Q |
| B-OPGRS-L6 | V | I | V | Q | G | M | A | Q |
| B-OPGRS-L7 | I | L | V | Q | G | M | A | Q |
| B-OPGRS-L8 | T | L | V | Q | G | M | A | Q |
| B-OPGRS-L9 | I | L | V | Q | G | M | A | Q |
| B-OPGRS-L10 | M | L | V | Q | S | R | L | Q |
| SpecOPGRS-1 | A | V | V | P | S | L | A | E |
| SpecOPGRS-2 | T | V | H | Q | G | M | A | Q |
| SpecOPGRS-3 | G | T | V | A | S | L | A | E |
| SpecOPGRS-4 | G | V | Q | Q | S | T | A | E |
| SpecOPGRS-5 | G | T | V | A | S | L | A | E |
| SpecOPGRS-7 | T | L | V | E | S | T | L | L |
| SpecOPGRS-9 | T | V | A | Q | G | M | A | Q |
| SpecOPGRS-10 | G | T | V | A | S | L | A | E |
| SpecOPGRS-12 | T | V | T | Q | G | M | A | Q |
| PolyT1RS-2 | L | L | V | Q | G | M | A | Q |
| PolyT1RS-3 | I | L | V | Q | G | M | A | Q |
| PolyT1RS-4 | I | L | V | Q | G | M | A | Q |
| PolyT1RS-5 | I | L | V | Q | G | M | A | Q |
| PolyT1RS-6 | I | L | V | Q | G | M | A | Q |
| PolyT1RS-7 | V | L | V | Q | G | M | A | Q |
| PolyT1RS-9 | I | L | V | Q | G | M | A | Q |
| PolyT1RS-10 | I | L | V | Q | G | M | A | Q |
| PolyT1RS-11 | I | L | V | Q | G | M | A | Q |
| PolyT1RS-12 | T | L | V | Q | G | M | A | Q |
| PolyT2RS-2 | L | V | V | Q | G | M | A | Q |
| PolyT2RS-3 | I | L | V | Q | G | M | A | Q |
| PolyT2RS-4 | V | V | V | Q | G | M | A | Q |
| PolyT2RS-5 | V | V | I | Q | G | M | A | Q |
| PolyT2RS-6 | I | V | V | Q | G | M | A | Q |
| PolyT2RS-8 | I | L | V | Q | G | M | A | Q |
| PolyT2RS-9 | V | V | I | Q | G | M | A | Q |
| PolyT2RS-10 | V | V | V | Q | G | M | A | Q |
| PolyT2RS-11 | L | V | V | Q | G | M | A | Q |

Supplementary Table 10. Sequences of all EcLeuRS variants isolated via FACS for individual ncAA, specificity, and polyspecificity sorts.

| **LeuRS Position** | **M40** | **L41** | **T252** | **S496** | **Y499** | **Y527** | **H537** | **Notes** |
|---|---|---|---|---|---|---|---|---|
| *Codon* | *VNK* | *VNK* | *GCT* | *RST* | *NNY* | *NNT* | *NNT* | |
| A-LysN3RS-1 | G | P | T | T | C | Y | G | |
| A-LysN3RS-2 | L | P | T | T | I | G | G | |
| A-LysN3RS-3 | A | G | T | S | G | N | C | |
| A-LysN3RS-1.4 | G | P | T | T | L | G | F | |
| A-LysN3RS-1.6 | A | N | T | S | N | G | F | |
| A-LysN3RS-1.7 | G | T | T | G | T | T | G | |
| A-LysN3RS-1.8 | G | E | T | T | C | Y | G | |
| A-ATyrRS-1 | L | T | T | A | H | S | G | |
| A-ATyrRS-2 | L | T | T | A | H | S | G | |
| A-ATyrRS-3 | L | T | T | A | H | S | G | |
| A-ATyrRS-5 | L | T | T | A | H | S | G | |
| A-ATyrRS-9 | L | T | T | A | G | T | G | |
| A-ATyrRS-12 | Q | G | T | S | Y | Y | H | |
| SpecOPGRS-6 | L | T | A | A | T | T | G | |
| SpecOPGRS-8 | L | T | A | A | C | C | G | |
| SpecOPGRS-11 | L | T | A | A | T | T | G | |
| PolyT1RS-8 | L | A | A | A | S | D | S | |
| PolyT2RS-7 | G | E | A | T | H | F | G | Q2K mutation |
| PolyT2RS-12 | G | A | A | A | S | F | A | |
| B-BockRS-1 | P | A | A | G | G | I | G | |
| B-BockRS-2 | A | G | A | S | G | T | G | |
| B-BockRS-3 | G | A | A | A | C | I | G | |
| B-BockRS-4 | A | G | A | S | G | T | G | |
| B-BockRS-5 | A | G | A | S | G | T | G | |
| B-BockRS-6 | A | G | A | S | G | T | G | |
| B-BockRS-7 | A | G | A | S | G | T | G | |
| B-BockRS-8 | A | G | A | S | G | T | G | |
| B-BockRS-9 | G | G | A | T | T | F | G | |
| B-BockRS-10 | P | G | A | G | A | V | G | |
| B-BockRS-11 | P | G | A | S | L | T | G | |
| B-BockRS-12 | A | A | A | G | A | C | G | |
| B-LysAlkRS-1 | A | G | A | S | V | I | G | |
| B-LysAlkRS-2 | S | P | A | G | T | V | G | |
| B-LysAlkRS-3 | M | H | A | G | A | G | G | |
| B-LysAlkRS-4 | G | V | A | G | T | V | L | |
| B-LysAlkRS-5 | P | G | A | G | C | C | G | |
| B-LysAlkRS-6 | A | G | A | S | G | T | G | |
| B-LysAlkRS-7 | A | G | A | S | G | T | G | |
| B-LysAlkRS-8 | A | G | A | S | G | T | G | |
| B-LysAlkRS-9 | A | G | A | S | G | T | G | |
| B-LysAlkRS-10 | M | L | A | G | F | S | G | |
| B-LysAlkRS-11 | A | G | A | S | G | T | G | |
| B-LysAlkRS-12 | A | G | A | S | G | T | G | |
| B-LysN3RS-1 | G | P | A | G | A | C | G | |
| B-LysN3RS-2 | P | G | A | G | A | V | G | |
| B-LysN3RS-3 | P | G | A | G | A | C | G | |
| B-LysN3RS-4 | P | G | A | G | C | C | G | |
| B-LysN3RS-5 | P | G | A | G | C | C | G | |
| B-LysN3RS-6 | A | P | A | G | G | H | G | |
| B-LysN3RS-7 | S | P | A | G | I | H | G | |
| B-LysN3RS-8 | A | P | A | G | G | H | G | |
| B-LysN3RS-9 | A | G | A | S | G | T | G | |
| B-LysN3RS-10 | A | G | A | S | G | T | G | |
| B-LysN3RS-11 | S | P | A | G | I | H | G | |
| B-LysN3RS-12 | P | G | A | G | C | C | G | |

Supplementary Table 11. Expected peptide sizes for the tryptic digest fragment containing the H54TAG codon from the scFv-Fc form of Donkey 1.1. Both cAA misincorporation and ncAA incorporation are included. Note that serine is the WT residue in the Donkey1.1 reporter that does not contain a TAG codon at H54.

| **Expected peptide fragment (H54 peptide only)** | | | | |
|---|---|---|---|---|
| | *Residue* | *MW (Da)* | *Diff. from WT (Da)* | *Peptide Mass (Da)* |
| **cAAs** | *Serine* | 105.1 | 0 | 2234.1 |
| | *Alanine* | 89.1 | -16 | 2218.1 |
| | *Arginine* | 174.2 | 69.1 | 2303.2 |
| | *Asparagine* | 132.1 | 27 | 2261.1 |
| | *Aspartic acid* | 133.1 | 28 | 2262.1 |
| | *Cysteine* | 121.2 | 16.1 | 2250.2 |
| | *Glutamic acid* | 147.1 | 42 | 2276.1 |
| | *Glutamine* | 146.2 | 41.1 | 2275.2 |
| | *Glycine* | 75.1 | -30 | 2204.1 |
| | *Histidine* | 155.2 | 50.1 | 2284.2 |
| | *Isoleucine* | 131.2 | 26.1 | 2260.2 |
| | *Leucine* | 131.2 | 26.1 | 2260.2 |
| | *Lysine* | 146.2 | 41.1 | 2275.2 |
| | *Methionine* | 149.2 | 44.1 | 2278.2 |
| | *Phenylalanine* | 165.2 | 60.1 | 2294.2 |
| | *Proline* | 115.1 | 10 | 2244.1 |
| | *Threonine* | 119.1 | 14 | 2248.1 |
| | *Tryptophan* | 204.2 | 99.1 | 2333.2 |
| | *Tyrosine* | 181.2 | 76.1 | 2310.2 |
| | *Valine* | 117.2 | 12.1 | 2246.2 |
| **ncAAs** | *OmeY* | 195.2 | 90.1 | 2324.2 |
| | *OPG* | 219.2 | 114.1 | 2348.2 |
| | *AcF* | 207.2 | 102.1 | 2336.2 |
| | *AzF* | 206.2 | 101.1 | 2335.2 |
| | *AzMF* | 220.2 | 115.1 | 2349.2 |
| | *IPhe* | 291.1 | 186 | 2420.1 |
| | *BPhe* | 209 | 103.9 | 2338 |
| | *DOPA* | 197.2 | 92.1 | 2326.2 |
| | *ATyr* | 196.2 | 91.1 | 2325.2 |
| | *LysN3* | 259.3 | 154.2 | 2388.3 |
| | *LysAlk* | 228.2 | 123.1 | 2357.2 |
| | *BocK* | 246.3 | 141.2 | 2375.3 |
| | *APhe* | 180.2 | 75.1 | 2309.2 |

Supplementary Table 12. Sequences of DOPARS clone variants isolated following error-prone mutagenesis and FACS. Note that silent mutations (one or more base pair mutations that do not cause a change in residue at that codon) are not included on this table.

| Clone | Q18 | D21 | T37 | L49 | K59 | Q63 | V108 | A109 | F111 | N132 | M179 | C185 | A186 | K195 | I209 | T231 | G242 | T263 | A264 |
|---|---|---|---|---|---|---|---|---|---|---|---|---|---|---|---|---|---|---|---|
| DOPARS-0.1-1 | - | - | - | S | - | - | - | - | - | - | - | - | - | - | - | - | - | - | - |
| DOPARS-0.1-2 | - | - | G | - | - | - | - | - | L | - | N | - | - | N | - | - | - | - | - |
| DOPARS-0.1-3 | - | - | - | - | - | - | - | - | - | - | - | - | - | - | - | I | - | - | - |
| DOPARS-0.1-5 | - | - | - | - | - | - | - | - | - | - | - | - | T | - | - | - | D | - | - |
| DOPARS-0.1-6 | - | - | - | - | R | R | - | - | - | - | - | - | - | - | - | - | - | - | - |
| DOPARS-0.1-7 | - | - | - | - | R | R | - | - | - | - | - | - | - | - | - | - | - | - | - |
| DOPARS-0.1-9 | - | A | - | - | - | - | - | - | - | - | - | Y | - | - | - | - | - | - | Q |
| DOPARS-0.1-10 | - | - | - | - | - | - | - | - | - | - | - | R | - | - | V | - | - | - | - |
| DOPARS-0.1-11 | - | - | - | - | - | - | - | V | - | - | - | - | - | - | - | - | - | - | - |
| DOPARS-0.1-12 | - | - | - | - | R | - | - | - | L | - | - | - | - | - | - | - | - | - | - |
| DOPARS-1-2 | - | - | - | - | R | - | - | - | - | - | - | - | - | - | - | - | - | - | T |
| DOPARS-1-3 | - | - | - | - | - | - | - | - | L | - | - | - | - | - | - | - | - | - | - |
| DOPARS-1-4 | - | - | - | - | - | - | - | - | L | - | - | - | - | - | - | - | - | - | - |
| DOPARS-1-5 | - | - | - | - | - | - | I | - | - | - | - | - | - | - | - | - | - | - | - |
| DOPARS-1-7 | - | - | - | - | - | - | - | - | - | S | - | - | V | - | - | - | - | - | - |
| DOPARS-1-8 | - | - | - | - | - | - | - | - | - | - | - | R | - | - | - | - | - | - | - |
| DOPARS-1-9 | - | - | - | - | - | - | - | - | - | - | - | - | - | - | - | - | - | A | - |
| DOPARS-1-10 | - | - | - | - | - | - | - | - | - | - | - | - | V | - | V | - | - | - | - |
| DOPARS-1-11 | - | - | - | - | - | - | - | - | - | - | - | - | T | - | - | - | - | - | - |
| DOPARS-1-12 | R | - | - | - | - | - | - | - | - | - | - | - | - | - | - | - | - | A | - |

Supplementary Table 12 continued

| Clone | F277 | M278 | I280 | S293 | Q300 | K321 | M353 | M360 | Q369 | K377 | T378 | N382 | E389 | K390 | S392 | Y396 | K415 |
|---|---|---|---|---|---|---|---|---|---|---|---|---|---|---|---|---|---|
| DOPARS-0.1-1 | - | - | - | - | - | - | - | - | - | - | - | - | - | - | - | - | - |
| DOPARS-0.1-2 | - | - | - | - | R | - | - | - | - | - | - | - | - | - | - | - | - |
| DOPARS-0.1-3 | - | - | - | - | - | - | - | - | - | - | - | - | - | - | - | - | - |
| DOPARS-0.1-5 | - | - | - | - | - | - | - | - | - | - | - | - | - | - | P | - | N |
| DOPARS-0.1-6 | - | - | - | - | - | - | - | - | - | - | - | - | - | - | - | - | - |
| DOPARS-0.1-7 | - | - | - | - | - | - | - | - | - | - | - | - | - | - | - | - | - |
| DOPARS-0.1-9 | - | - | T | - | - | - | - | - | - | E | - | - | D | - | - | - | - |
| DOPARS-0.1-10 | - | - | - | - | - | - | V | - | - | - | - | - | - | - | - | H | - |
| DOPARS-0.1-11 | - | - | - | - | - | R | - | - | - | - | - | - | - | - | - | - | - |
| DOPARS-0.1-12 | - | - | - | G | - | - | - | - | - | - | - | - | - | - | - | - | - |
| DOPARS-1-2 | - | - | - | - | - | - | - | - | - | - | - | - | - | - | - | - | - |
| DOPARS-1-3 | - | V | - | - | - | - | V | - | - | - | A | S | - | - | - | - | - |
| DOPARS-1-4 | - | - | - | - | - | - | - | - | - | - | - | - | - | - | - | - | - |
| DOPARS-1-5 | L | - | - | - | - | - | - | - | - | - | - | - | - | - | - | - | - |
| DOPARS-1-7 | - | - | - | - | - | - | - | T | R | - | - | - | - | - | - | - | - |
| DOPARS-1-8 | - | - | - | - | - | - | - | - | | - | - | - | - | - | - | - | - |
| DOPARS-1-9 | - | - | - | - | - | - | - | - | - | - | - | - | - | R | - | - | - |
| DOPARS-1-10 | - | - | - | - | - | - | - | - | - | - | - | - | - | - | - | - | - |
| DOPARS-1-11 | - | - | - | - | - | - | - | - | - | - | - | - | - | - | - | - | - |
| DOPARS-1-12 | - | - | - | - | - | - | - | - | - | - | - | - | - | R | - | - | - |

Supplementary Table 13. Flow cytometry primary and secondary labeling conditions and reagents.

| Detection | Primary label (dilution) | Secondary label (dilution) |
|---|---|---|
| HA tag (displaying) | Mouse anti-HA (1:500) | Goat anti-mouse Alexa Fluor 488 (1:500) |
| c-Myc tag (full length) | Chicken anti-CMYC (1:500) | Goat anti-chicken Alexa Fluor 647 (1:500) |
| Biotin | None | Streptavidin Alexa Fluor 488 (1:500) |
